# Recovering the Impacts of Landscape Changes through the Demographic History of a Western Ghats Endemic Bird

**DOI:** 10.64898/2026.11.24.768910

**Authors:** T R Rayis, Chiti Arvind, Ian Lockwood, Ashwin Warudkar, Naman Goyal, Meghana Natesh, V V Robin

## Abstract

Global landscapes are shaped by long-term palaeoclimatic changes and more recent, rapid anthropogenic changes. Changes in the demographic history of a species correspond to responses to changes in its natural habitat. Recent developments in genomics allow the reconstruction of recent historical effective population sizes. Yet only a limited number of studies have linked these changes to specific land-use changes in their natural habitat, especially with known trajectories of habitat change. The Western Ghats mountaintops - Shola Sky Islands have been modified through a combination of colonial-era changes, more recent spread of agriculture and invasive woodlands. In these cloud forests, we selected four characteristic landscape change situations: a) habitat decrease, b) habitat increase (from a spread of woody invasives), c) a combination of decrease followed by an increase (woody invasives), and d) no change. We assessed how these landscape changes affect the demographic history of an endemic passerine, the White-bellied Sholakili (*Sholicola albiventris*). We generated whole genome sequences for 84 individuals across six populations, each with contrasting land-use changes, and assessed their historical effective population sizes (Ne) estimated over the last 100 generations using Linkage Disequilibrium (LD) methods (*GONE2*). We used historical landscape information from published documents, historical maps, satellite imagery, and archived photographs to correlate land-use change information with changes in Ne. Further, we assessed these findings using genomic simulations. We found a clear correlation between the demographic histories of White-bellied Sholakili populations and landscape changes in Shola sky islands over the last century. In one case, our simulations suggest possible artefacts from population fusion. We also recognised a previously unknown population expansion in the species, underscoring the importance of genomic tools and simulations for understanding population dynamics and informing assessments of conservation actions such as restoration.

## 1. Introduction

Global landscapes are continuously changing, shaped by both long-term paleoclimatic and more recent human-induced alterations (Alverson et al., 2013). Paleoclimate-driven changes, for example, glacial-interglacial cycle-driven changes in forest and grassland habitats (Ray et al., 2001; Servant et al., 1993; van der Hammen, 1974), sea-level change-driven changes in freshwater and terrestrial ecosystem connectedness (Voris, 2000), monsoon intensity-driven greening and aridification in the Sahara (Kowalski et al., 1989; Larrasoaña et al., 2013; Petit-Maire & Guo, 1997) have repeatedly altered the connectivity, distribution, and structure of ecosystems worldwide. These changes influence ecosystem functioning, species distribution, and global species diversity (Breeze et al., 2016; Hora, 1953; Svenning et al., 2015). While paleoclimatic changes have long-term effects on ecosystems, recent human-induced changes are much more rapid and may therefore have a greater impact, as many species are unable to respond to rapid changes (Quintero & Wiens, 2013; C. D. Thomas et al., 2004). On a global scale, anthropogenic changes primarily convert forests, grasslands, wetlands, and other native landscapes into human settlements, agricultural fields, grazing grounds, and commercial plantations. Deforestation in the Amazon rainforests (Foley et al., 2007), land conversion in Southeast Asian forests (Sodhi et al., 2010), and woody invasions in the savanna and grassland ecosystems worldwide (Stevens et al., 2017) are examples of such rapid changes in the last few decades. Studies worldwide indicate such changes may have a direct or indirect impact on the resident species.

The responses of resident species to human-induced landscape changes are complex and vary globally. Across the globe, examples such as native fauna of the Amazon rainforest (Decaëns et al., 2018), grazers of Tanzanian savannas (Kimaro et al., 2019), and grassland-dependent birds of Western Ghats grasslands (Ramesh et al., 2025) suggest a widely known negative impact of anthropogenic changes on native fauna. However, a lesser-known potential positive impact on resident species, mainly due to their adaptability to the change, is documented in cases such as browsers of Tanzanian savannas (Kimaro et al., 2019), Golden Jackals across Europe and Asia (Stefanović et al., 2024), and some generalist bird species in India (Viswanathan et al., 2024). Thus, despite being rapid, the impacts of recent landscape changes are highly species and landscape-dependent and hence need to be carefully assessed before making inferences for conservation or restoration.

Across the wide variety of topographic and climatic zones in the tropics (Beck et al., 2023; Jain et al., 2022), the impacts of anthropogenic changes are evident. India, being one of the major hubs of such landscape changes, has documentation of these changes, majorly dating back to the colonial times (Joshi et al., 2018; Tian et al., 2014). The montane cloud forests in the Western Ghats in India, specifically the Shola Sky Islands, are well documented, owing to their use in colonial times as retreats and summer capitals for their wet, cool habitats (Joshi et al., 2018). This colonial usage also resulted in landscape changes of the native forest grassland mosaic (Robin & Nandini, 2012), over the past two centuries (Arasumani et al., 2018, 2019a; Ramesh et al., 2025). Across the Sky Islands, the type and extent of landscape changes vary with factors such as agriculture, human settlements, and commercial plantations, including tea and cardamom, that replace native forests and grasslands (Arasumani et al., 2018, 2019a; Ramesh et al., 2025). The colonial usage is also marked by a recent increase in woodland cover, driven by invasion of woody timber species, mainly into the grasslands (Arasumani et al., 2019b; Ramesh et al., 2025). Consequently, in some parts of the Sky Islands, a sequence of woodland decrease and increase has occurred (Arasumani et al., 2019). However, some areas in the west, which are part of protected areas such as Eravikulam National Park, have remained wetter and relatively stable, with few changes (Arasumani et al., 2019a; Rice, 1984). These varying changes across the landscape are known to have different impacts on the resident species. While (Ramesh et al., 2025) found a recent decline in grassland-dependent bird species following landscape changes, (Jobin et al., 2025) found that forest-dependent bird species showed no significant preference for natural landscapes over human-modified wooded habitats. Moreover, (Arunima et al., 2026) reported an overall increase in landscape connectivity for forest-specialist birds in the sky islands, whereas grassland-specialist birds showed a decrease. (Natesh et al., n.d.) (in prep.) found that, historically (10kya to 200kya), endemic birds restricted to the Shola sky islands showed less fluctuation and more stability in effective population size (Ne) than widespread birds, indicating its importance as a climatic refugia, similar to other tropical mountain tops across the world (He et al., 2016; Hewitt, 2000; MacVean & Schuster, 1981; Stewart et al., 2010). This indicates the importance of analysing the impact of recent rapid human-induced changes on these endemic species. White-bellied Sholakili (*Sholicola albiventris*) (WBS) (Figure 1) is a forest specialist passerine endemic to the Shola Sky Islands of the Palani-Anamalai hill range (Robin et al., 2006; Robin & Sukumar, 2002). The species has multiple genetically varying populations across its habitat (Arvind et al., 2025), which are susceptible to fragmentation, habitat loss (Sreekumar & Nameer, 2022), and varying land-use changes across its range (Arasumani et al., 2019a) (Figure 1), impacting its connectivity and occupancy (Arunima et al., 2026; Jobin et al., 2025). The Shola sky islands, with their contrasting land-use changes, and the White-bellied Sholakili, with its variable responses to these changes, offer an interesting study system for analysing connections between landscape changes and the resulting genomic signatures in resident populations.

**Figure 1:**
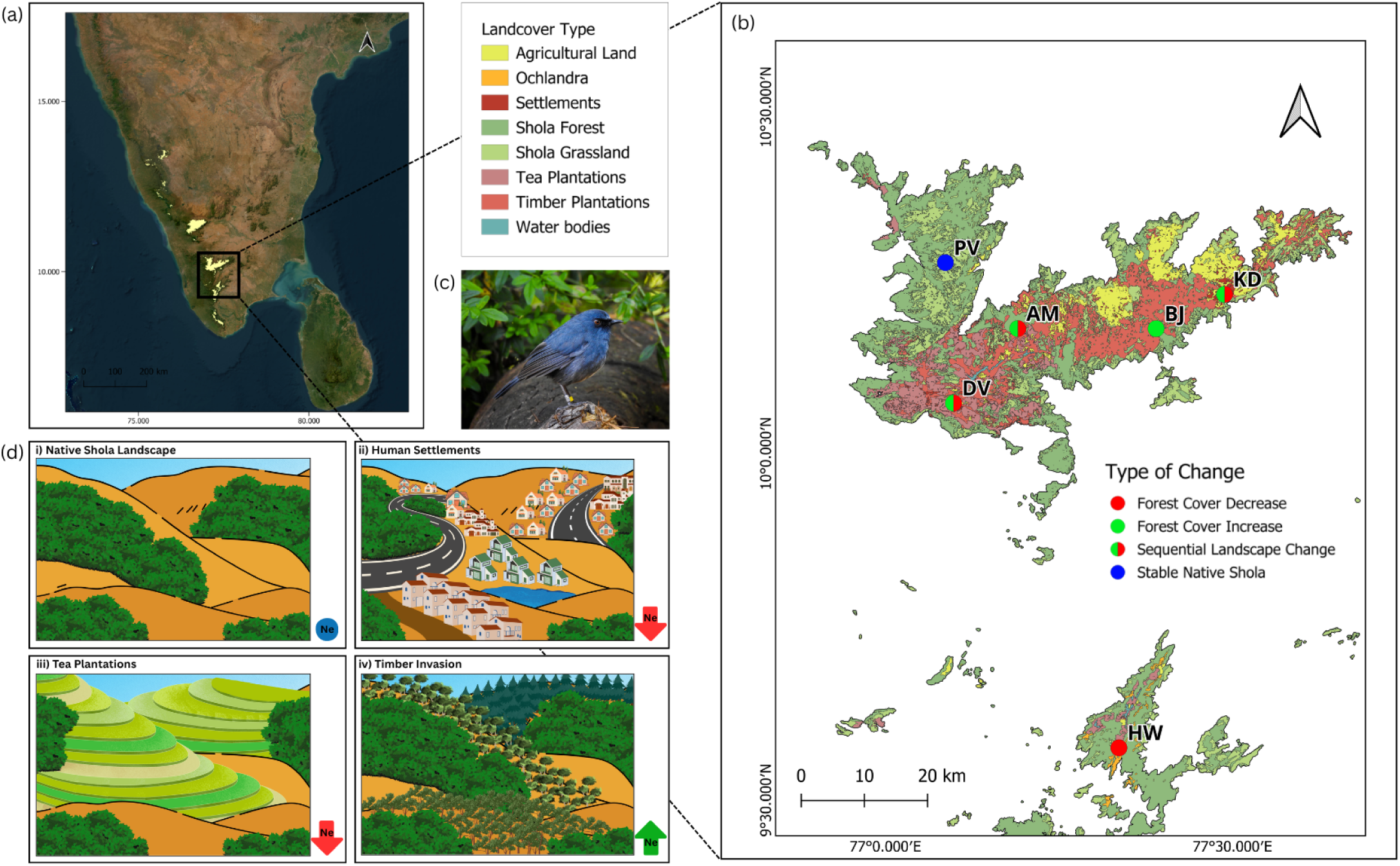
The study system. (a) Peninsular India with the Shola sky islands (>1400m in elevation) highlighted. (b) Map of Anamalai-Palani sky islands with landcover types and six sampling locations. HW - HighWavies, BJ - Berijam, KD - Kodaikanal, PV - Poovar. Map redrawn from (Arasumani et al., 2019b) (c) Study species - White-bellied Sholakili (Sholicola albiventris; WBS) (Photo credits: Vinay KL) (d) Types of landcover changes and the expected change in effective population size (Ne) of WBS. i) Native Shola landscape with a mosaic of forests (dark green) and grasslands (yellow-brown). Ne expected to be stable. ii&iii) Forest cover reduction by Human settlements and Tea plantations, respectively. Ne expected to reduce in both cases. iv) Forest cover increase by Invasive Timber species. Ne expected to increase. (Illustrations made in Canva).

Recent developments in genomics present multiple tools for estimating the demographic history or the change in historic effective population size (Ne) (Wright, 1931) of a population or species of interest (Excoffier et al., 2021; Li & Durbin, 2011; Liu & Fu, 2015; Santiago et al., 2020; Schiffels & Durbin, 2014; Wang et al., 2016). The estimated demographic history can indicate how landscape changes impact the species and its adaptability to the natural environment. Several studies worldwide have linked historical changes in Ne to landscape changes and climatic fluctuations (Kumar et al., 2025; Mays et al., 2018; Nadachowska-Brzyska et al., 2015; Natesh et al., 2020; Pujolar et al., 2022). Most such Ne estimation methods, which rely on fixed mutation, lack the ability to infer Ne over more recent timescales (Santiago et al., 2020). However, this limitation was overcome by recent developments in tools that use Linkage Disequilibrium (LD) methods to infer past Ne (Barbato et al., 2015; Hayes et al., 2003; Santiago et al., 2020). *GONE2 (Santiago et al., 2025)* uses LD-based analyses to estimate changes in Ne over the last 150 generations in populations of interest, thereby providing information on the impacts of recent human-induced changes on those populations. The recent demographic history estimates from such tools have been effectively utilized to monitor species dynamics and identify populations of conservation importance (Khan et al., 2024; Kyriazis et al., 2025; Quinn et al., 2024; von Seth et al., 2022). However, only a limited number of studies have linked these demographic trends to specific recent landscape changes within a natural, comparative framework (Benham et al., 2025; Stefanović et al., 2024). The ability of such methods to capture changes across populations that have undergone varying levels of landscape change, as in the Shola sky islands, remains unclear. Moreover, it remains unclear whether the inferences from such studies are truly biological or technical artefacts arising from population structure that affect LD patterns (Novo et al., 2023). We try to answer these research gaps using the Sky Island model.

We generated whole-genome sequences from 84 individuals of White-bellied Sholakili across six populations, each with distinct, contrasting landscape changes over the last 200 years (Figure 1; Table 1). We use LD-based analyses, historic landscape information, and genomic simulations to answer two main questions:

1. Do we recover changes in demographic history (Ne) that correspond with changes in the landscape following three specific scenarios

a. Populations with contrasting changes to their landscape - increase vs decrease in forest cover
b. Populations that have gone through sequential changes in the same landscape (decrease followed by increase)
c. Population with no significant change to its native landscape
2. Can we use genomic simulations to examine whether the patterns observed are truly biological or a technical artefact of the data?

a. Is the recent Ne expansion driven by an actual population increase or by population fusion (two or more previously isolated populations contribute equally to the formation of a new population)?
b. Is the recent Ne decline driven by an actual population decrease or by population fragmentation?

**Table 1:** Populations of WBS selected for the study with the number of samples, the observed landscape changes in the location, and the expected trends in Ne for each population.

| **Population<br>(Number of samples)** | **Landscape Change Type** | **Expectation of Ne change** |
|---|---|---|
| HighWavies: HW (13) | Tea plantations from the 1940s. No invasion or increase in habitat | A recent decline, with no increase |
| Berijam: BJ (11) | Timber invasion into the grasslands from the 1970s | A recent increase, with no decline |
| Kodaikanal: KD (15) | Human settlements from 1845 to the early 1900s; Timber invasion from the 1970s | An initial decline followed by a recent increase |
| Devikulam: DV (15) | Tea plantations from the 1870s to 1940; Timber invasion from 1970s | An initial decline followed by a recent increase |
| Anamudi: AM (20) | Timber from the 1970s; Land conversion by the local community in the 1880s | An initial decline followed by a recent increase |
| Poovar (Eravikulam National Park): PV (10) | Stable native shola; no invasion, no habitat loss | No change |

## 2. Methods

### 2.1 Study design

#### 2.1.1 Study site selection

The study sites, and thereby the populations of White-bellied Sholakili, were selected based on historical landscape change data from Arasumani et al. (2019), field observations, and sample availability. Accordingly, six populations (HighWavies (HW), Berijam (BJ), Kodaikanal (KD), Devikulam (DV), Anamudi (AM), and Poovar (PV) (Table 1)) were selected, with three different landscape change histories:

##### a: Contrasting landscape changes

Impacts of contrasting landscape changes were tested in the HW and BJ populations. HW experienced a reduction in forest cover starting around 1946, due to extensive timber plantations established by clearing a large portion of the forest (Hutton & David, 2008). On the other hand, a recent increase in invasive timber woodlands in BJ is observed from 1973 to 2017, with almost 60% of grasslands lost to timber invasion, while forests remained largely unchanged <u>(Arasumani et al. 2019)</u>.

##### b: Sequential landscape changes

Impacts of sequential landscape changes were tested in the KD, DV, and AM populations. Transformation of native forests and grasslands to human settlements and agriculture began in the 1840s and has continued to expand over the years (Arasumani et al., 2019a; Mitchell, 1972). DV had a major tea plantation drive in the region from the 1870s to 1940, some of which involved clearing native forest (Suresh et al., 2018). On the other hand, in AM, a potential forest cover reduction from land use by the native tribal community was noted by (Munro, 1880). Moreover, <u>(Arasumani et al. 2019)</u> identified a net increase in forest cover across all three regions from 1973 to 2017 due to invasive timber.

##### c: No landscape change

PV, as part of the Eravikulam National Park, remained an undisturbed native shola over the years, with minimal human intervention (Rice, 1984) and was selected as a control population.

#### 2.1.5 Validating Ne estimations

Genomic simulations with a baseline, population fusion, and population fragmentation (see section 2.4 for further details) were used to validate historic Ne estimates.

### 2.2 Genomic data generation

#### 2.2.1 Sample selection

Sampling for all populations was conducted between 2021 and 2023 using mist-netting (Arvind, 2025; Warudkar, 2025), following the method described by (Robin et al., 2010). A drop of blood was collected from the birds’ ulnar vein. The collected blood was then stored using Queen’s Lysis Buffer (QLB) and FTA cards. The blood samples in QLB were frozen until further extractions. A total of 84 samples were selected across the landscape, with the constraint that each population should have at least nine individual samples, following (Khan et al., 2024) (Table 1).

#### 2.2.2 DNA Extraction and Sequencing

DNA was extracted from the selected samples using the Qiagen DNeasy Blood and Tissue Kit, with minor modifications to the manufacturer’s protocol (Vinay, n.d.). The concentrations of the extracted DNA were measured using the Qubit 4 fluorometer (Thermo Fisher Scientific, Inc.) and Qubit 1X dsDNA HS Assay Kit (Thermo Fisher Scientific, Inc.) following the manufacturer’s protocol. Library Preparation and DNA Sequencing of the samples were outsourced to Neuberg Diagnostics Pvt. Ltd. Library preparation was carried out using the KAPA HyperPlus Kit; libraries were quantified using a Qubit 4 fluorometer, and library insert size was assessed using a TapeStation 4150 (Agilent). The whole genomes were sequenced, targeting ∼15X coverage on the Illumina NovaSeq 6000 platform (150 bp, paired-end).

#### 2.2.3 Genomic Data Preprocessing

From the raw reads, adapter sequences and low-quality reads were then trimmed out using *Trimmomatic v0.39* (Bolger et al., 2014). The raw and trimmed reads were inspected for sequence quality, GC content, and adapter contamination using *FastQC v0.12.1* (*Babraham Bioinformatics - FastQC A Quality Control Tool for High Throughput Sequence Data*, n.d.). The trimmed paired reads were then mapped to the White-bellied Sholakili scaffold-level reference assembly from NCBI genomes (GCA_050043525.1; Scaffold N50: 68.6 Mb) (Vinay et al., 2025) using *bwa v0.7.18-r1243* with the mem option (Li, 2013), and read group information were added. The SAM output was then converted to BAM format and sorted using *samtools v1.20* (Li et al., 2009). The mapped raw BAMs were then filtered for unmapped reads and low-quality mapping using *samtools*. The filtered BAMs were then resorted using *samtools*. The PCR duplicates were identified and removed using *Picard Tools v2.27.5 MarkDuplicates* (*Picard*, n.d.), with metrics recorded for the samples. The final BAMs were indexed using *samtools*. The BAMs was assessed using *Qualimap v2.3 bamqc* (Okonechnikov et al., 2016) to evaluate genomic coverage and mapping statistics for each of the raw, filtered, and PCR-duplicate-removed BAMs.

#### 2.2.4 Variant calling and filtering

Further analyses were conducted at the population level, repeating the downstream steps for each of the six populations. The variant sites from the final BAMs were identified independently using both *bcftools v1.10.2* (Li, 2012) and *freebayes v1.3.6* (Garrison & Marth, 2012). To obtain only the high-confidence sites, only the variant sites recognised by both bcftools and freebayes were retained using *bedtools v2.31.1* (Quinlan & Hall, 2010). The intersected VCF was compressed and indexed for downstream processing. The intersected VCF was then filtered to retain only biallelic SNPs using *bcftools v2.31.1* (Danecek et al., 2021), and indels were removed using *vcftools v0.1.16* (Danecek et al., 2011). Additional stringent filtering was done using *vcftools* to retain only the high-quality variant sites. The small scaffolds (<20Mb) and the sex linked scaffolds (scaffold33 and scaffold35 - identified using the approach in Natesh et al (in prep.)) were removed from the filtered VCF, retaining only the large scaffolds (scaffolds1 - scaffold14). The final filtered VCFs were converted to PED and MAP formats using *PLINK v1.90b7.2* (Purcell et al., 2007). For analyses other than demographic estimations, regions in Linkage Disequilibrium (LD) were identified with the *PLINK* sliding-window approach and pruned with *vcftools*.

### 2.3 Demographic analysis

#### 2.3.1 Assessment of landscape change across two centuries from six sites

Historical landscape data for the Anamalai-Palani hill ranges were obtained from multiple sources, including published historical documents, archived landscape photographs, and historical maps. Published literature was used to obtain temporal information on some landscapes prior to the 1970s (Hutton & David, 2008; Lekshmi, 2020; Mitchell, 1972; Munro, 1880; Suresh et al., 2018). Archived historical landscape photographs of KD and BJ, captured by Dr Nora Mitchell, were acquired with the help of Ian Lockwood. These images were used to provide visual evidence of landscape changes in these regions from the 1960s. Historical maps based on satellite imagery, published by Arasumani et al. (2019), were used to obtain landscape information post-1970s. Additionally, historical maps by Lockwood et al. (in prep.), based on Survey of India (SOI) topographic sheets, were used to obtain landscape classification data for the study sites from the 1920s, except for HighWavies. Together, these sources were used to obtain temporal information and sequences of landscape changes across the study sites in Anamalai-Palani hill ranges. A buffer of 5 km radius was created around the six sampling points in the maps of Lockwood et al. (in prep.) and (Arasumani et al., 2019a) in *QGIS 3.40.15*. The percentages of each land cover type were calculated to analyse quantitative changes in the landscape across three timelines, i.e., the 1920s, 1973, and 2017 .

#### 2.3.2 Testing assumptions of panmixia within populations

To ensure that the assumption of panmixia required for linkage-disequilibrium-based demography estimates (Novo et al., 2023; Waples & England, 2011) is not violated, pairwise relatedness and population-structure tests were conducted for each population. Pairwise Relatedness among individuals in each population was estimated using *ngsRelate v2.0* (Hanghøj et al., 2019). To remove closely related individuals from the demographic analysis, one individual from pairs with pairwise relatedness (rab) greater than 0.1 was removed from the downstream analysis. Population structure within each population was assessed using *ADMIXTURE v1.3.0* (Alexander et al., 2009) to ensure the absence of substructure. Analyses were conducted for five values of K (1-5), and cross-validation (CV) errors were calculated for each. The K value with the smallest CV error value was identified as the most likely number of genetic clusters. To further evaluate population structure, principal component analysis (PCA) was performed in *PLINK*. The eigenvectors and eigenvalues from *PLINK PCA* were visualised to examine any further clustering patterns in the dataset. The same approaches were repeated for a VCF containing all six populations to assess variation across all the individuals.

#### 2.3.3 Estimating the recombination rate of White-bellied Sholakili

Recombination rates required for demographic inferences in *GONE2* were estimated with *iSMC* (Barroso et al., 2019). For the analysis, the individual with the highest average sequencing depth was selected from the KD population. The analysis was restricted to include only scaffolds larger than 100 kb, as shorter scaffolds provide limited coalescent-based information. The *bamCaller.py* scripts from *msmc-tools* pipeline (Schiffels & Wang, 2020) were used to do variant calling and mask generation. Variants were called independently for each scaffold using *bcftools mpileup* and *call* functions with stringent filters. Masks were generated with *bamCaller.py,* applying a depth filter based on the individual’s calculated mean depth. The VCFs and corresponding mask files identifying high-confidence callable regions were generated as outputs. The VCF files generated for each scaffold were concatenated and sorted. A tab-separated file specifying scaffold IDs and the start and end coordinates was generated using the *generate_tab.py* scripts from the *iSMC* pipeline. The mask files from all scaffolds were merged and sorted to generate a genome-wide mask file. A “negative mask” complementary to the generated mask was prepared using *bedtools* subtract. This mask was further trimmed to include only the regions present in the tab file. iSMC was run with parameter optimisation, posterior decoding, and sequence summarising. The output rho (population-scaled recombination rate; ρ) estimates were averaged in windows of 10kb, 100kb, and 1 Mb using the ismc_mapper. The Ne value for the KD population was calculated from Watterson’s theta (Θ_w_), estimated using a modified version of *genomics_vcf_summary_stats.py* (Funk, n.d.). The rho value was converted to recombination rate (r) in cM/Mb using the following formula.

*p* = *4*Ne*r* ; r = recombination rate in crossovers per base pair per generation *r_cMMb* = *r*10e8* ; r_cMMb = recombination rate in cM/Mb

*Θ_w_* = *4*Ne*μ* ; μ = mutation rate

#### 2.3.4 Variation in demographic histories of populations that have undergone different scenarios of land-use changes

Historic Ne for the past 150 generations were estimated for each population using *GONE2* (Santiago et al., 2025). A constant recombination rate of 1 cM/Mb (-r 1) from the rho estimates was used with all other default parameters. The PED and MAP files generated using *PLINK* from the non-LD-pruned VCF of panmictic individuals were used as inputs. The program was run for 100 bootstrap iterations, and the mean Ne for each time point was calculated. Results for generations older than the 100th generation were excluded, following (Khan et al., 2024). The generation time of 3.18 years reported by (Bird et al., 2020) and the sampling year were used to convert generations to years in the dataset. The results were visualised together with temporal information from the historical landscape data.

*Calendar years = Sampling year - generations ago * generation time*

### 2.4 Validating demographic history estimations using genomic simulations

Information from the demographic history patterns of four populations (HW, BJ, KD, and PV - selected based on differing patterns of demography) was used as input for *SLiM v5.0* (Haller et al., 2025) to run genomic simulations. All the simulation runs were initialised with nucleotide-based and multi-chromosome models, using 14 chromosomes, each of a size similar to the first 14 scaffolds of the White-bellied Sholakili reference. The simulations utilised only genomic elements with neutral nucleotide-based mutations, with a species-specific mutation rate of 7.31e-9 (estimated using the generation time (3.18 years) from (Bird et al., 2020) and the mutation rate for Collard Flycatchers (CF) (2.3e-9) from (Smeds et al., 2016)), implemented via a Jukes-Cantor model matrix. The default recombination rate of 1e-8 crossovers per base pair per generation (1 cM/Mb) was used in the simulations. Each simulation was run for an initial burn-in of 10000 ticks (generations) followed by another 1000 ticks, during which tweaks to subpopulation sizes and numbers were applied as needed. At the end of the last tick (11000th tick), chromosome-level VCFs of randomly selected 20 individuals from the final population/s were generated as output. These VCFs were combined into a single VCF using *bcftools concat*, filtered to remove invariant sites, and indexed. The final VCF was converted to PED and MAP formats using *PLINK,* then used to run *GONE2* with the same parameters as in the demographic analysis. The results were visualised alongside the real data and simulation input to compare and validate the observations from real populations.

For each of the four populations, two sets of simulations were run: one with a real increase or decrease in Ne (hereafter, the ‘baseline model’), and the other with population fusion or fragmentation, to assess potential artefacts in the dataset. We ran population fusion models in populations with Ne expansion (with populations merging at the point of Ne increase) and population fragmentation models in populations with Ne decline (with populations splitting at the point of Ne decrease). The models for the BJ and PV populations were run at a smaller scale than the original estimates, while keeping the patterns consistent. To validate the recent Ne increase pattern in the KD and BJ populations, two sets of simulations were run: first, the baseline model with an actual population increase and second, a population fusion model. For the baseline model, a single subpopulation in SLiM was simulated, with its population size varied from approximately the 10950th generation to match one of the KD or BJ bootstraps. For the population fusion model, 2 and 4 subpopulations (based on the number of large shola forest patches in the sampling location) were simulated in the KD and BJ-based models, respectively, from the 10000th generation onwards. The population size was varied to match the true population until its point of increase, at which all subpopulations were combined into a single subpopulation. To further explore the possibility of real population expansion in the KD population, GONE2 with the subpopulation assumption (-x flag) was run. To validate the recent Ne decline pattern in HW and KD populations, two sets of simulations were run: first, the baseline model, similar to previous but with an actual population decline and second, a population fragmentation model. For the population fragmentation model, a single population was simulated to undergo changes similar to those in real populations until its decline, at which point a subpopulation split was simulated, forming two subpopulations. Results from initial runs were used to make tweaks and rerun the SLiM models until a satisfactory level of similarity to the real population was achieved.

## 3. Results

### 3.1 Preliminary Quality Check to ensure data quality

A total of 84 samples were sequenced (Table S1). All individual samples had an average coverage of 10X or higher after PCR duplicate removal, except for one sample from the KD population (Table S1), which was therefore excluded from the analysis.

### 3.2 Demographic analysis

#### 3.2.1 Contrasting and sequential landscape changes across two centuries in six sites

The quantified bar plots of land cover types across three time points revealed the extent of landscape changes in all six regions post-1920s (except for HW) (Figure S1). The analysis recovered a clear signal of an increase in forest cover by invasive woodlands across multiple locations. Moreover, the archived historical photographs (Figure S1b) from BJ and KD further validated the presence of timber plantations in the region dating back to the 1960s. Various published documents provided timelines for forest cover reduction events across multiple populations, most of which predate the 1920s (see Section 2.1.1).

#### 3.2.2 Detection of violation in panmictic assumptions in some populations

The relatedness analysis (*ngsRelate)* (Figure S2) indicated that some individuals within the HW, KD, DV and AM populations exhibit significant pairwise relatedness (rab) values.

The population structure analysis *(ADMIXTURE)* (Figure S3) indicated no clear substructure within each population, as the cross-validation (CV) error is smallest for K = 1 across all populations. However, marginal substructure with increasing K values can be observed in some populations. This is expected, as some selected individuals from these populations show significant pairwise relatedness. Some unrelated individuals in AM and PV also showed clustering at higher K values.

The principal component analysis (PCA) results across multiple PCs (Figures S4, S5, and S6) indicate patterns similar to the structure plots. No significant clustering is observed in any population except the PV population; a single individual separates from the rest of the population cluster on the major PCs.

Based on these results and the fraction of missing data, one individual from the HW population, three from the KD, two from DV, six from AM and one from PV were excluded from the demographic analysis to avoid violation of the panmixia assumption (Novo et al., 2023; Waples & England, 2011). The final numbers of individuals for the demographic analyses ranged from 9 - 14 in these 6 populations (Table S1).

#### 3.2.3 Estimated recombination rates fall outside the known recombination rates of birds

The estimated rho values (population-scaled recombination rates) ranged from 1.92e-5 (0.085 cM/Mb) to 0.253 (1120 cM/Mb), with a positively skewed distribution (mean = 0.045 (199 cM/Mb); median = 0.023 (101 cM/Mb)) (Figure S8). The high positive skewness is most likely an artefact of the dataset, since the calculated recombination rates are unrealistic for birds (Backström et al., 2010; Bascón-Cardozo et al., 2024; Kawakami et al., 2014). The most frequently observed rho value was 0.00053 (2.34 cM/Mb), which falls inside the known recombination rates for birds. The dataset showed another minor peak at 5.74e-5 (0.25 cM/Mb), which also falls within the plausible recombination rates for birds. Since the dataset showed a high variation in the rho estimate, a recombination rate of 1cM/Mb, which is between the two observed peaks, was taken as a proxy for the GONE2 analyses and genomic simulations.

The Watterson’s Theta (Θ_w_) estimated for the KD population was 1.65e-4, corresponding to an Ne of 5642.95, which was used to convert the rho values to recombination rates in cM/Mb.

#### 3.2.4 Variation in recent demographic history across six populations of White-bellied Sholakili

The GONE2 estimates of the HW and BJ populations revealed the impact of contrasting landscape changes in WBS. The HW population showed a gradual decline from the past, followed by a much steeper decline from the 1970s onward (Figure 2a1). The tea plantations in HW were established in the 1940s (Hutton & David, 2008), resulting in a major conversion of the landscape to tea plantations. While the start of the Ne decline didn’t coincide with the start of tea plantations in the area, the decline occurred within two decades. The BJ population showed an earlier Ne decline (1840s) followed by a much bigger Ne expansion (almost 100-fold) towards the 1940s (Figure 2a2). The large Ne expansion aligned with the timeline of timber invasion in BJ, during which most of the region’s grasslands were converted to timber woodlands (Arasumani et al., 2019a).

**Figure 2:**
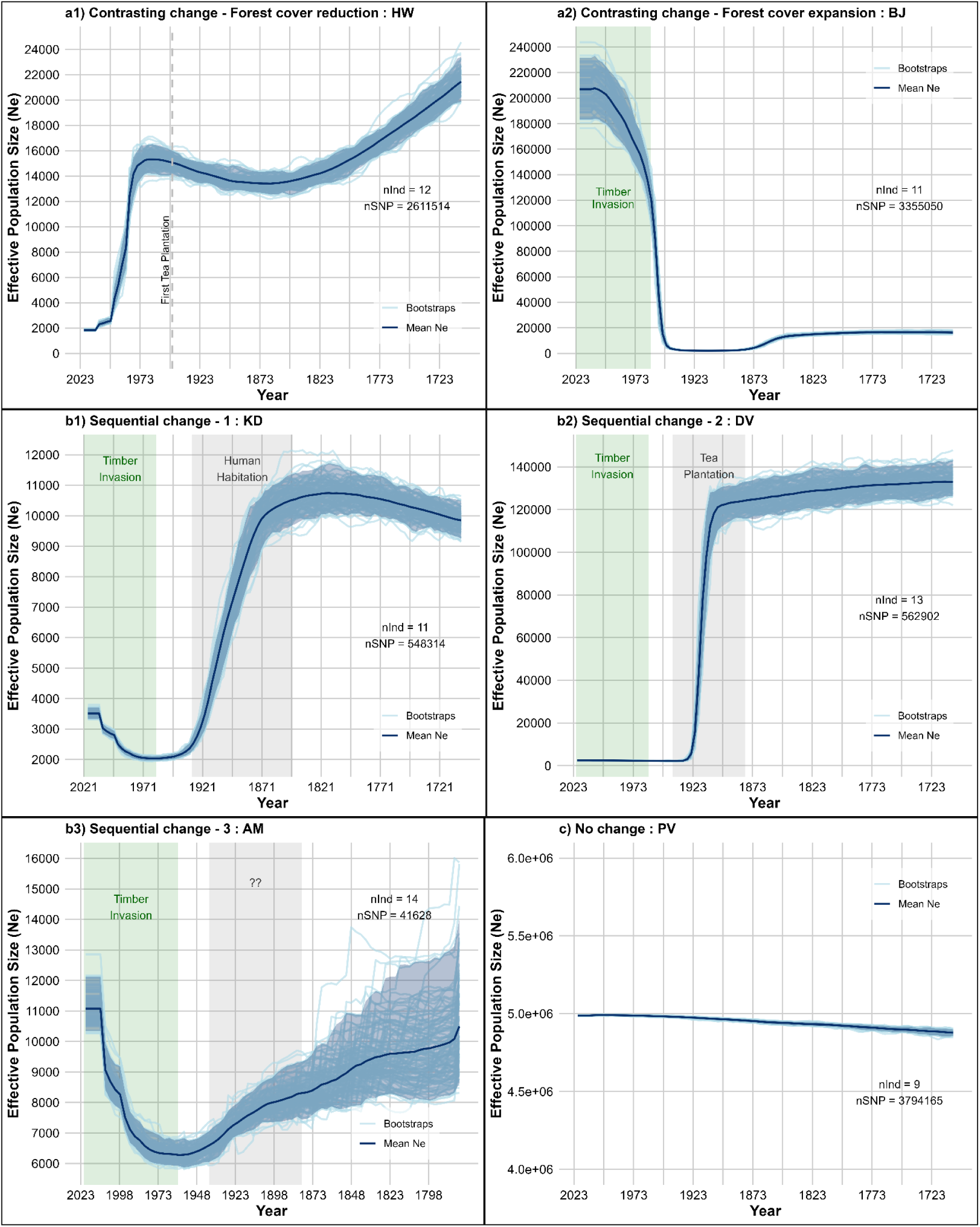
Recent demographic history of each population estimated by GONE2 for the past 100 generations (scaled to years using generation time = 3.18): The x-axis represents years back in time, the origin being the sampling year. The y-axis is the effective population size (Ne). Note the different scales of the y-axis in each plot. The light blue lines represent each independent bootstrap result, and the dark blue line represents the mean of all bootstraps. The dark blue shaded region represents the 95% Confidence Interval (CI). The number of individuals (nInd) and the number of SNPs (nSNP) in each of the analyses are given inside each plot. a) Demographic history of populations with contrasting landscape changes: a1) HW population with forest cover reduction. The dotted lines represent the start of forest clearing for tea plantations. a2) BJ population with forest cover increase. The light green shade is the time scale during which invasive forest cover expansion occurred. b) Demographic historis of populations with sequential landscape changes. b1) KD population. b2) DV population b3) AM populations. The region shaded in grey is the known timescale of human settlements and tea plantations in Kodaikanal and Devikulam, respectively. The light green-shaded region indicates the same as in a2. c) Demographic history of the PV population with no landscape change. The y-axis is scaled from 4e6 to 6e6.

The estimates from KD, DV, and AM recovered the demographic histories in response to sequential landscape changes. The KD population showed a historic Ne decline beginning around the 1840s, followed by a gradual increase towards the more recent years (1970s) (Figure 2b1). The DV (Figure 2b2) and AM (Figure 2b3) populations showed a similar trend, but the population decline events happened about 20 years later than in KD. Even though the timing of population increase was the same across all three populations, the extent of the increase differed: KD showed the highest fold change, followed by AM, while DV had only a slight increase in Ne. The timing of these population change events coincided with known historical timelines of landscape change. The Ne decline in KD from the 1840s corresponded with the start of human settlements in KD (Mitchell, 1972), and the subsequent steep decline coincided with the construction of major buildings, roads, and Kodaikanal Lake. On the other hand, the decline in DV and AM temporally matched with the tea plantation drives from the 1870s to 1940 (Suresh et al., 2018) and the conversion of the forest by the native human community (Munro, 1880), respectively. The gradual increase in Ne over the following years in all three populations coincided with a major invasion of non-native woodland (Arasumani et al., 2019a).

The estimates from the PV population showed a relatively stable but very high Ne (Figure 2c), following the unaltered habitat.

### 3.3 Genomic simulations

#### 3.3.1 Potential impact of population fusion on recent population increase

The baseline model of HW populations (Figure S10) captured the decline in Ne consistent with the input, although the timing of the decline was shifted later than expected (by almost 5 generations). In further simulations, the timing of the estimated Ne decline matched the estimates from the HW population when the Ne drop was simulated further back in time (Figure 3a).

**Figure 3:**
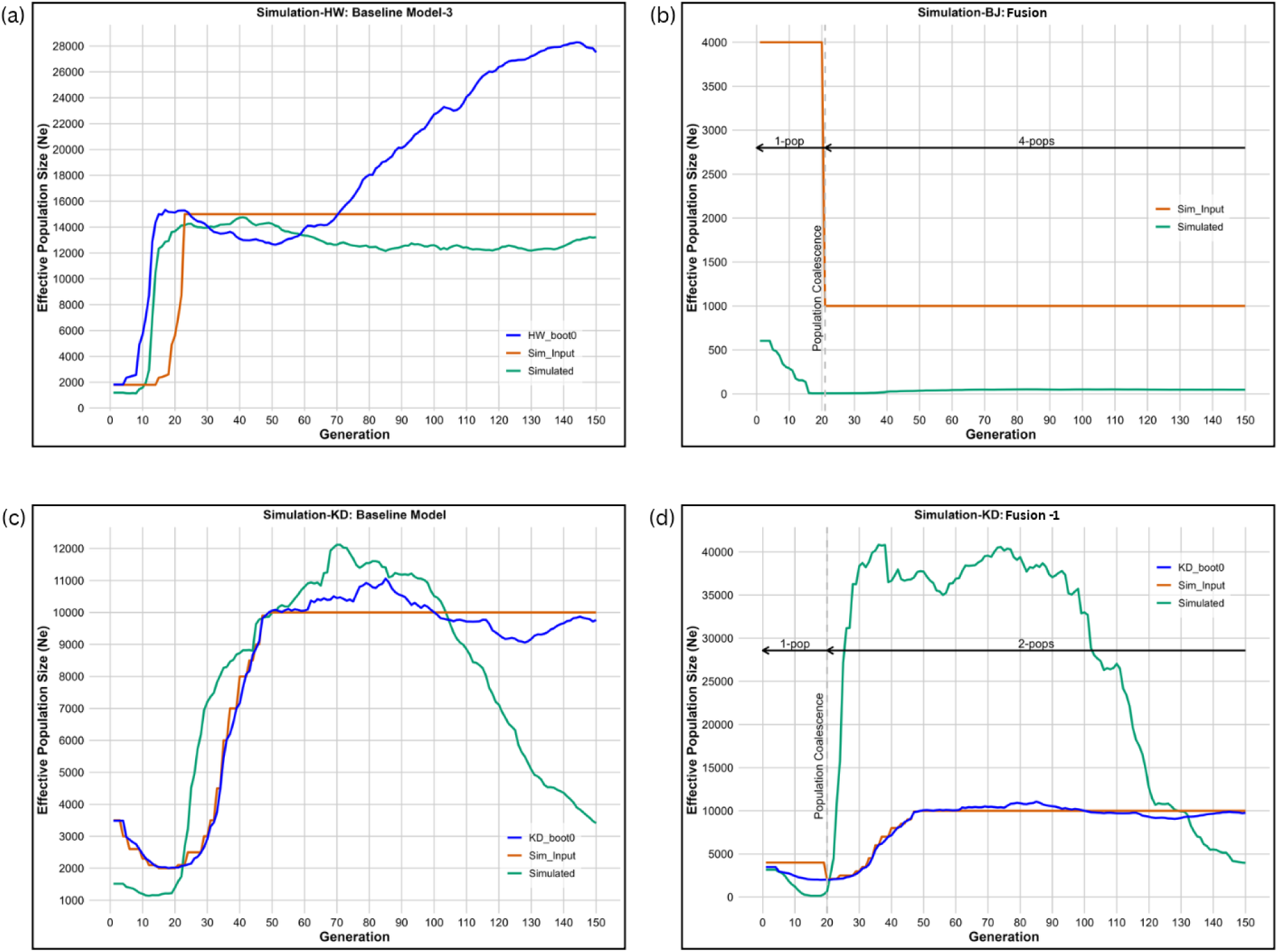
SLiM Simulation results. The dark blue line is the first GONE2 bootstraps (boot0) from each population. The orange line is the input given in SLiMgui. In fusion and fragmentation models, the orange line is the input population size of a single subpopulation. The green line is the output of GONE2 from 20 individuals randomly selected from ‘Generation 0’ of the simulated dataset. (a) Baseline model from HW simulations. (b) Fusion model from BJ simulations. (c & d) Baseline and Fusion models for KD simulations. The rest of the simulation results are in the supplementary.

The Ne estimated from the baseline model of BJ failed to capture the almost 100-fold increase in the input population (Figure S11). There was more than a twofold difference between the estimated Ne of the final simulated population (at generation 0) and the input value. Similarly, in KD simulations, there was a discrepancy between the simulation inputs and the Ne estimates from the simulated data. While the decline in Ne was captured well, the magnitude of the recent increase didn’t match the input values (Figure 3c). An almost twofold difference was observed in the final population, similar to that observed in BJ. Additionally, the estimated Ne for generations above 100 deviates considerably from the expected trajectory. The PV simulations (Figure S13) also showed variation from the input of a constant 1000 Ne, with the estimated Ne varying by ±300 from the inputs up to 100 generations. As with KD, beyond the 100th generation, the difference between the input and the estimates was much larger.

The fusion models captured the recent Ne expansion much better than the baseline models. The BJ fusion model (Figure 3b; S11), with four populations connected at the 20th generation to form one, effectively captured an almost 100-fold increase in Ne. However, the absolute values of the Ne estimates were much lower than the input values (Figure 3b; S11), probably due to the disparity between census population size and Ne. The KD fusion models have recovered the recent Ne expansion, but the estimated population decline was always steeper than expected, with consistently underestimated Ne values at the point of fusion (Figure 3d; S12). In both BJ and KD, the fusion event resulted in a decline in Ne at the point of fusion. Furthermore, the result from GONE2 estimates with a subpopulation assumption in KD revealed a very recent increase in the last 5 generations (Figure S9).

The fragmentation models failed to capture the population declines at the exact times. In the HW fragmentation model (Figure S10), no decline in estimated Ne was observed when the fragmentation event was in the last 11 generations. Similarly, in KD fragmentation models (Figure S12), the Ne estimates didn’t show considerable variation from the input when the fragmentation event occurred in the 20th generation. When fragmentation was simulated much earlier, at the 45th generation, the estimated Ne showed a very steep population decline of almost 10-fold over the last 10 generations, which did not align with the timestamps of the simulation inputs.

## 4. Discussion

### 4.1 The demographic history patterns for sampled populations closely reflect landscape changes

The study found that the impact of contrasting and sequential changes to the native landscape was well captured in the reconstructed demographic trajectory of a range-restricted endemic bird. The results indicated a decline in Ne in certain populations, temporally associated with a reduction in forest cover due to the conversion of native forest to human settlements, agriculture, and tea plantations. This observation is similar to many other human-made landscape modifications that negatively impact native fauna (Bourgeois et al., 2024; Decaëns et al., 2018; Fetene et al., 2019; Ramesh et al., 2025; Sirami et al., 2010). Some of these impacts have been documented through genomics (Ahrens et al., 2026; Arana et al., 2023; Benham et al., 2025; Bourgeois et al., 2024; Qin et al., 2024). However, apart from the decline, our study also revealed a recent Ne expansion across multiple populations of the species, coinciding with an increase in woody invasives in these regions. This observation suggests a possible unique positive impact of human interventions on the species, driven by the expansion of forest cover, with woodland invasion replacing native grasslands. This inference points to either a new adaptation (Friis et al., 2022; Lo Cascio Sætre et al., 2017; Mascarenhas et al., 2023) or niche tracking (Avidad et al., 2025; Ramirez et al., 2025; Tingley et al., 2009) in the species, allowing it to expand into invasive timber woodlands. Similar observations for White-bellied Sholakili were also made using ecological data in the occupancy study by (Jobin et al., 2025) and the connectivity study by (Arunima et al., 2026). A recent population increase, due to the species’ adaptability, niche tracking, or successful restoration efforts, has also been documented in a few other studies using genomic methods (Ahrens et al., 2026; Benham et al., 2025; Stefanović et al., 2024; Teng et al., 2025). Our study identified a similar, previously unidentified trend in a range-restricted, endemic specialist bird, closely linked to changes in the native landscape. A unique feature of our study was that we were able to track alternating population trajectories at a very fine scale - a decline, followed by an increase. Such trajectories might be useful for examining other anthropogenic impacts and subsequent restoration.

### 4.2 Genomic simulations suggest a possible population fusion

Our simulation results helped disentangle true population-size changes from potential artefacts of population fusion and fragmentation in the inferred Ne trajectory. The results supported population decline in certain populations of the species. However, our results also suggested a lag of approximately 5-10 generations before the population decline was detected as a genomic signal. Although the Ne decline signal was aptly captured, we found that a possible artefact of population fusion may be driving the recent Ne expansion signals in some populations. In certain other populations, the patterns were driven by both an actual population increase and an artefact of population fusion, as suggested by demographic estimates that assume population substructure. A similar expansion signal in estimated Ne, driven by both a known population size expansion and the fusion of previously isolated populations, was observed in Eurasian otters (*Lutra lutra*) in the UK (N. E. Thomas et al., 2025). Moreover, the reduced ability of LD-based analysis to accurately estimate Ne beyond 100 generations was also observed. This contrasts with some earlier papers, which have drawn inferences from generations beyond 100 (Stefanović et al., 2024; Teng et al., 2025). Among the many studies that have used genomic tools to estimate historical Ne, only a few have explored the possibility that the patterns are driven by artefacts of population structure (Benham et al., 2025; Khan et al., 2024; Li & Durbin, 2011; Novo et al., 2023; Pérez-Sorribes et al., 2024; Sharma et al., 2012). Our study indicates the need for caution before drawing general inferences from LD-based Ne estimates. Even though our study suggests an artefact of recent population fusion in the Ne expansion of a forest-specialist endemic bird, ecologically it signals population connectivity due to the expansion of woody invasives into the surrounding grassland, and the species’ ability to effectively inhabit and move within them. A general observation on specialist endemic birds worldwide is the negative impact of human-induced landscape changes, primarily due to habitat fragmentation (Carrara et al., 2015; de Lima et al., 2013; Ferrer-Sánchez & Rodríguez-Estrella, 2015). Here, this study suggests that when changes increase habitat connectivity for such species (Arunima et al., 2026), the impact on their demographic history can be positive. The study thus indicates that the effects of these changes are highly landscape- and species-specific, depending on each species’ ability to utilise the newly formed habitat.

The estimated Ne for some of the population is unusually high. Even though we did not have measures to connect the Ne to the actual population size (Nc) for White-bellied Sholakili at the time of the study, field observations suggest that this is one of the most common understory birds in the Shola landscape. However, we advise caution when drawing inferences from the absolute values of Ne in our results; these patterns are best interpreted in comparison with the different landscape features. Further, we recognise the lack of species-specific recombination rate (default 1 cM/Mb, which falls in the known avian recombination range is used (Backström et al., 2010; Groenen et al., 2009; Kawakami et al., 2014; Stapley et al., 2017)) and the absence of a chromosome-level assembly for WBS that could improve our estimates.

### 4.3 Conclusion

Using the Shola sky island system, we found a close link between land-use changes and the recent demographic history of an endemic forest-specialist bird. Our results suggest that the demographic response of resident species to habitat alteration is context-dependent, which may vary across landscapes and species. We thus underline the importance of species-specific approaches in conservation efforts, as not all landscape changes impact species negatively, and changes that benefit one species may have the opposite effect on another. The study also underscores the importance of genomic tools and techniques for monitoring species demography and the need to validate them before making inferences. The framework presented in the study is also replicable in areas with active restoration efforts worldwide, to monitor whether a resident species or population of interest is recovering. The study thus provides an example of using genomics and simulations to answer questions that inform conservation measures or monitoring of restoration efforts.

## Supporting information

Supplementary methods & results

## Ethics

All animal handling was done with permission from the Kerala Forest Department and the Tamil Nadu Forest Department.

## Data Accessibility

All code used in this study will be available on request.

## Declaration of AI use

Grammarly was used to check grammar and improve sentence structure. ChatGPT 5 and Claude (Sonnet 5) were used to debug scripts written by the authors. All scientific content, analyses, interpretations, and conclusions are the authors’ own work. The authors take full responsibility for the accuracy and integrity of this work.

## Conflict of interest declaration

The authors declare no competing interests

## Funding

This work was supported by Anusandhan National Research Foundation (ANRF) CRG/2022/001182 and Rohini Nilekani Philanthropies grant G-202410-00940 to VVR.

## Acknowledgements

We thank the Tamil Nadu Forest Department and Kerala Forest department for granting the necessary permits to conduct this study. We are grateful to the field team for their help with blood sample collection. We thank Dr Anubhab Khan and Dr Enrique Santiago for their guidance with the demographic history analysis. We thank Vishwa Jagati, Dr Praveen P, Gayathri Ramesh, Archita Sharma and members of the Bird Lab at the Indian Institute of Science Education and Research, Tirupati, for feedback on the project. We also thank the Kodaikanal International School for providing accommodation during the fieldwork.

## Notes

### Competing Interest Statement

The authors have declared no competing interest.

## References

1. Ahrens, C. W., Miller, A. D., Silver, L. W., McLennan, E. A., Hogg, C. J., & Weeks, A. R. (2026). Escaping bottlenecks: The demographic path to genetic recovery in koalas (Phascolarctos cinereus). *Science (New York*, N.Y.), 391(6789), 1010–1014.

2. Arana, A., Esteves, J., Ramírez, R., Galetti, P. M., Jr, Pérez Z, J., & Ramirez, J. L. (2023). Population genomics reveals how 5 ka of human occupancy led the Lima leaf-toed gecko (Phyllodactylus sentosus) to the brink of extinction. Scientific Reports, 13(1), 18465.

3. Arasumani, M., Khan, D., Das, A., Lockwood, I., Stewart, R., Kiran, R. A., Muthukumar, M., Bunyan, M., & Robin, V. V. (2018). Not seeing the grass for the trees: Timber plantations and agriculture shrink tropical montane grassland by two-thirds over four decades in the Palani Hills, a Western Ghats Sky Island. PloS One, 13(1), e0190003.

4. Arasumani, M., Khan, D., Vishnudas, C. K., Muthukumar, M., Bunyan, M., & Robin, V. V. (2019a). Invasion compounds an ecosystem-wide loss to afforestation in the tropical grasslands of the Shola Sky Islands. Biological Conservation, 230, 141–150.

5. Arasumani, M., Khan, D., Vishnudas, C. K., Muthukumar, M., Bunyan, M., & Robin, V. V. (2019b). Invasion compounds an ecosystem-wide loss to afforestation in the tropical grasslands of the Shola Sky Islands. Biological Conservation, 230, 141–150.

6. Arunima, J., Arvind, C., Jobin, V., Lele, A., & Robin, V. V. (2026). Changes in habitat connectivity for range-restricted birds reflect patterns of woodland invasion. Biological Conservation, 317(111800), 111800.

7. Arvind, C. (2025). Population and individual level variation in birdsong. Indian Institute of Science Education and Research Tirupati.

8. Arvind, C., Sawant, S., Srinivasan, R., Arpitha, G. P., Goyal, N., Warudkar, A., & Robin, V. V. (2025). Distinct, asynchronous processes underlie cultural and genetic variation in a bird species with complex song repertoires. In bioRxiv. 10.1101/2025.11.02.684965

9. Avidad, P. M., Clavero, M., & Viana, D. S. (2025). Climate and land-use change leading to niche expansion and shifts in birds. Journal of Biogeography, 52(2), 443–453.

10. Babraham Bioinformatics - FastQC A Quality Control tool for High Throughput Sequence Data. (n.d.). Retrieved November 18, 2025, from https://www.bioinformatics.babraham.ac.uk/projects/fastqc/

11. Backström, N., Forstmeier, W., Schielzeth, H., Mellenius, H., Nam, K., Bolund, E., Webster, M. T., Ost, T., Schneider, M., Kempenaers, B., & Ellegren, H. (2010). The recombination landscape of the zebra finch Taeniopygia guttata genome. Genome Research, 20(4), 485–495.

12. Barbato, M., Orozco-terWengel, P., Tapio, M., & Bruford, M. W. (2015). SNeP: a tool to estimate trends in recent effective population size trajectories using genome-wide SNP data. Frontiers in Genetics, 6, 109.

13. Barroso, G., V., Puzović, N., & Dutheil, J. Y. (2019). Inference of recombination maps from a single pair of genomes and its application to ancient samples. PLoS Genetics, 15(11), e1008449.

14. Bascón-Cardozo, K., Bours, A., Manthey, G., Durieux, G., Dutheil, J. Y., Pruisscher, P., Odenthal-Hesse, L., & Liedvogel, M. (2024). Fine-scale map reveals highly variable recombination rates associated with genomic features in the Eurasian blackcap. Genome Biology and Evolution, 16(1). 10.1093/gbe/evad233

15. Beck, H. E., McVicar, T. R., Vergopolan, N., Berg, A., Lutsko, N. J., Dufour, A., Zeng, Z., Jiang, X., van Dijk, A. I. J. M., & Miralles, D. G. (2023). High-resolution (1 km) Köppen-Geiger maps for 1901–2099 based on constrained CMIP6 projections. Scientific Data, 10(1), 724.

16. Benham, P. M., Cicero, C., Davila, M. M., Enbody, E. D., Miller, K. S., Shultz, A. J., Smith, L. L., Nachman, M. W., & Bowie, R. C. K. (2025). Patterns of genetic diversity within three California quail species are best explained by climate and landscape changes. Molecular Ecology, 34(20), e70093.

17. Bird, J. P., Martin, R., Akçakaya, H. R., Gilroy, J., Burfield, I. J., Garnett, S. T., Symes, A., Taylor, J., Şekercioğlu, Ç. H., & Butchart, S. H. M. (2020). Generation lengths of the world’s birds and their implications for extinction risk. Conservation Biology: The Journal of the Society for Conservation Biology, 34(5), 1252–1261.

18. Bolger, A. M., Lohse, M., & Usadel, B. (2014). Trimmomatic: a flexible trimmer for Illumina sequence data. Bioinformatics (Oxford, England), 30(15), 2114–2120.

19. Bourgeois, Y., Warren, B. H., & Augiron, S. (2024). The burden of anthropogenic changes and mutation load in a critically endangered harrier from the Reunion biodiversity hotspot, Circus maillardi. Molecular Ecology, 33(6), e17300.

20. Breeze, P. S., Groucutt, H. S., Drake, N. A., White, T. S., Jennings, R. P., & Petraglia, M. D. (2016). Palaeohydrological corridors for hominin dispersals in the Middle East ∼250–70,000 years ago. Quaternary Science Reviews, 144, 155–185.

21. Carrara, E., Arroyo-Rodríguez, V., Vega-Rivera, J. H., Schondube, J. E., de Freitas, S. M., & Fahrig, L. (2015). Impact of landscape composition and configuration on forest specialist and generalist bird species in the fragmented Lacandona rainforest, Mexico. Biological Conservation, 184, 117–126.

22. Danecek, P., Auton, A., Abecasis, G., Albers, C. A., Banks, E., DePristo, M. A., Handsaker, R. E., Lunter, G., Marth, G. T., Sherry, S. T., McVean, G., Durbin, R., & 1000 Genomes Project Analysis Group. (2011). The variant call format and VCFtools. Bioinformatics (Oxford, England), 27(15), 2156–2158.

23. Danecek, P., Bonfield, J. K., Liddle, J., Marshall, J., Ohan, V., Pollard, M. O., Whitwham, A., Keane, T., McCarthy, S. A., Davies, R. M., & Li, H. (2021). Twelve years of SAMtools and BCFtools. GigaScience, 10(2). 10.1093/gigascience/giab008

24. Decaëns, T., Martins, M. B., Feijoo, A., Oszwald, J., Dolédec, S., Mathieu, J., Arnaud de Sartre, X., Bonilla, D., Brown, G. G., Cuellar Criollo, Y. A., Dubs, F., Furtado, I. S., Gond, V., Gordillo, E., Le Clec’h, S., Marichal, R., Mitja, D., de Souza, I. M., Praxedes, C., … Lavelle, P. (2018). Biodiversity loss along a gradient of deforestation in Amazonian agricultural landscapes. Conservation Biology: The Journal of the Society for Conservation Biology, 32(6), 1380–1391.

25. de Lima, R. F., Dallimer, M., Atkinson, P. W., & Barlow, J. (2013). Biodiversity and land-use change: understanding the complex responses of an endemic-rich bird assemblage. Diversity & Distributions, 19(4), 411–422.

26. Excoffier, L., Marchi, N., Marques, D. A., Matthey-Doret, R., Gouy, A., & Sousa, V. C. (2021). Fastsimcoal2: Demographic inference under complex evolutionary scenarios. *Bioinformatics (Oxford*, England), 37(24), 4882–4885.

27. Ferrer-Sánchez, Y., & Rodríguez-Estrella, R. (2015). Man-made environments relationships with island raptors: endemics do not cope with habitat changes, the case of the island of Cuba. Biodiversity and Conservation, 24(2), 407–425.

28. Fetene, A., Yeshitela, K., & Gebremariam, E. (2019). The effects of anthropogenic landscape change on the abundance and habitat use of terrestrial large mammals of Nech Sar National Park. Environmental Systems Research, 8(1). 10.1186/s40068-019-0147-z

29. Friis, G., Atwell, J. W., Fudickar, A. M., Greives, T. J., Yeh, P. J., Price, T. D., Ketterson, E. D., & Milá, B. (2022). Rapid evolutionary divergence of a songbird population following recent colonization of an urban area. Molecular Ecology, 31(9), 2625–2643.

30. Funk, E. (n.d.). genomics_tools. Github. Retrieved March 20, 2026, from https://github.com/erikrfunk/genomics_tools.git

31. Garrison, E., & Marth, G. (2012). Haplotype-based variant detection from short-read sequencing. In arXiv [q-bio.GN]. arXiv. http://arxiv.org/abs/1207.3907

32. Groenen, M. A. M., Wahlberg, P., Foglio, M., Cheng, H. H., Megens, H.-J., Crooijmans, R. P. M. A., Besnier, F., Lathrop, M., Muir, W. M., Wong, G. K.-S., Gut, I., & Andersson, L. (2009). A high-density SNP-based linkage map of the chicken genome reveals sequence features correlated with recombination rate. Genome Research, 19(3), 510–519.

33. Haller, B. C., Ralph, P. L., & Messer, P. W. (2025). SLiM 5: Eco-evolutionary simulations across multiple chromosomes and full genomes. In bioRxivorg. 10.1101/2025.08.07.669155

34. Hanghøj, K., Moltke, I., Andersen, P. A., Manica, A., & Korneliussen, T. S. (2019). Fast and accurate relatedness estimation from high-throughput sequencing data in the presence of inbreeding. GigaScience, 8(5), giz034.

35. Hayes, B. J., Visscher, P. M., McPartlan, H. C., & Goddard, M. E. (2003). Novel multilocus measure of linkage disequilibrium to estimate past effective population size. Genome Research, 13(4), 635–643.

36. He, K., Hu, N.-Q., Chen, X., Li, J.-T., & Jiang, X.-L. (2016). Interglacial refugia preserved high genetic diversity of the Chinese mole shrew in the mountains of southwest China. Heredity, 116(1), 23–32.

37. Hewitt, G. (2000). The genetic legacy of the Quaternary ice ages. Nature, 405(6789), 907–913.

38. Hora, S. L. (1953). THE SATPURA HYPOTHESIS. Science Progress, 41(162), 245–255.

39. Hutton, A. F., & David, P. (2008). Note on a collection of snakes from south India, with emphasis on the snake fauna of the meghamalai hills (high wavy mountains). Journal of The Bombay Natural History Society, 105(3), 299–316.

40. Jain, S., Tiwari, V., Thapa, A., Mangla, R., Jaiswal, R. K., Kumar, V., Tiwari, S., Tulbure, M. G., Galkate, R., Lohani, A. K., & Pandey, K. (2022). Performance evaluation of Google earth engine based precipitation datasets under different climatic zones over India. Remote Sensing in Earth Systems Sciences, 5(4), 263–276.

41. Jobin, V., Harikrishnan, C. P., Lele, A., Joshi, V., Chanda, R., Lawrence, S., Aravind, P. S., Mubeen, M., Joseph, M. R., Subash, M., Nandini, R., Das, A., Jathanna, D., & Robin, V. V. (2025). Invasion blurs the biphasic mosaic, amplifies the “wicked problem” through impacts on avian distributions in montane habitats. In Ecology (No. Biorxiv;2025.04.03.646526v1). bioRxiv. https://www.biorxiv.org/content/10.1101/2025.04.03.646526v1.full.pdf

42. Joshi, A. A., Sankaran, M., & Ratnam, J. (2018). “Foresting” the grassland: Historical management legacies in forest-grassland mosaics in southern India, and lessons for the conservation of tropical grassy biomes. Biological Conservation, 224, 144–152.

43. Kawakami, T., Smeds, L., Backström, N., Husby, A., Qvarnström, A., Mugal, C. F., Olason, P., & Ellegren, H. (2014). A high-density linkage map enables a second-generation collared flycatcher genome assembly and reveals the patterns of avian recombination rate variation and chromosomal evolution. Molecular Ecology, 23(16), 4035–4058.

44. Khan, A., Sil, M., Thekaekara, T., Garg, K. M., Sinha, I., Khurana, R., Sukumar, R., & Ramakrishnan, U. (2024). Divergence and serial colonization shape genetic variation and define conservation units in Asian elephants. Current Biology: CB, 34(20), 4692–4703.e5.

45. Kimaro, H. S., Asenga, A. M., Munishi, L., & Treydte, A. C. (2019). Woody encroachment extent and its associated impacts on plant and herbivore species occurrence in Maswa Game Reserve, Tanzania. Environment and Natural Resources Research, 9(3), 63.

46. Kumar, H., Vijay, N., & Rajamani, N. (2025). Asian pika populations track local glaciation events through the Pleistocene. In bioRxiv. 10.1101/2025.06.25.661461

47. Kyriazis, C. C., Venkatraman, M., Masuda, B., Steiner, C. C., Cassin-Sackett, L., Crampton, L. H., Flanagan, A. M., Foster, J. T., Houck, M. L., Misuraca, A. C., Paxton, E. H., Robinson, J. A., Fleischer, R. C., Ryder, O. A., Campana, M. G., & Wilder, A. P. (2025). Population genomics of recovery and extinction in Hawaiian honeycreepers. Current Biology, 35(11), 2697–2708.e4.

48. Lekshmi, R. (2020). Management Plan of Anamudi Shola National Park 2020--21 to 2029--30. Kerala Forests and Wildlife Department. https://munnarwildlife.com/axFileDownload.php?dispname=Anamudi_Shola_National_Park_V10_Final_New.pdf&downName=uploads%2Fmanagementplan%2F3.pdf

49. Li, H. (2012). A statistical framework for SNP calling, mutation discovery, association mapping and population genetical parameter estimation from sequencing data. In arXiv [q-bio.GN]. arXiv. http://arxiv.org/abs/1203.6372

50. Li, H. 2013). Aligning sequence reads, clone sequences and assembly contigs with BWA-MEM. In arXiv [q-bio.GN]. arXiv. http://arxiv.org/abs/1303.3997

51. Li, H., & Durbin, R. (2011). Inference of human population history from individual whole-genome sequences. Nature, 475(7357), 493–496.

52. Li, H., Handsaker, B., Wysoker, A., Fennell, T., Ruan, J., Homer, N., Marth, G., Abecasis, G., Durbin, R., & 1000 Genome Project Data Processing Subgroup. (2009). The Sequence Alignment/Map format and SAMtools. Bioinformatics (Oxford, England), 25(16), 2078–2079.

53. Liu, X., & Fu, Y.-X. (2015). Exploring population size changes using SNP frequency spectra. Nature Genetics, 47(5), 555–559.

54. Lo Cascio Sætre, C., Coleiro, C., Austad, M., Gauci, M., Sætre, G.-P., Voje, K. L., & Eroukhmanoff, F. (2017). Rapid adaptive phenotypic change following colonization of a newly restored habitat. Nature Communications, 8(1), 14159.

55. MacVean, C., & Schuster, J. C. (1981). Altitudinal distribution of passalid beetles (Coleoptera, passalidae) and Pleistocene dispersal on the volcanic chain of northern central America. Biotropica, 13(1), 29.

56. Mascarenhas, R., Meirelles, P. M., & Batalha-Filho, H. (2023). Urbanization drives adaptive evolution in a Neotropical bird. Current Zoology, 69(5), 607–619.

57. Mays, H. L., Jr, Hung, C.-M., Shaner, P.-J., Denvir, J., Justice, M., Yang, S.-F., Roth, T. L., Oehler, D. A., Fan, J., Rekulapally, S., & Primerano, D. A. (2018). Genomic analysis of demographic history and ecological niche modeling in the endangered Sumatran rhinoceros Dicerorhinus sumatrensis. Current Biology, 28(1), 70–76.e4.

58. Mitchell, N. (1972). The Indian hill-station: Kodaikanal: No. 141. University of Chicago, Dept. of Geography.

59. Munro, J. D. (1880). The High Ranges of Travancore.

60. Nadachowska-Brzyska, K., Li, C., Smeds, L., Zhang, G., & Ellegren, H. (2015). Temporal dynamics of avian populations during Pleistocene revealed by whole-genome sequences. Current Biology, 25(10), 1375–1380.

61. Natesh, M., Vinay, K. L., Ghosh, S., Jayapal, R., Mukherjee, S., Vijay, N., & Robin, V. V. (2020). Contrasting trends of population size change for two Eurasian Owlet species—Athene brama and Glaucidium radiatum from south Asia over the late Quaternary. Frontiers in Ecology and Evolution, 8. 10.3389/fevo.2020.608339

62. Natesh, M., Vinay, K. L., Jagati, V., Goyal, N., Warudkar, A., Rayis, T. R., Vijay, N., & Robin, V. V. (n.d.). Demographic responses to historic climate changes are contingent upon ecological specialisation [Unpublished manuscript].

63. Novo, I., Ordás, P., Moraga, N., Santiago, E., Quesada, H., & Caballero, A. (2023). Impact of population structure in the estimation of recent historical effective population size by the software GONE. *Genetics, Selection*, Evolution: GSE, 55(1), 86.

64. Okonechnikov, K., Conesa, A., & García-Alcalde, F. (2016). Qualimap 2: advanced multi-sample quality control for high-throughput sequencing data. *Bioinformatics (Oxford*, England), 32(2), 292–294.

65. Pérez-Sorribes, L., Villar-Yanez, P., Smeds, L., & Mergeay, J. (2024). Comparing genetic Ne reconstructions over time with long-time wolf monitoring data in two populations. Evolutionary Applications, 17(10), e70022.

66. Picard. (n.d.). Retrieved November 18, 2025, from https://broadinstitute.github.io/picard/

67. Pujolar, J. M., Blom, M. P. K., Reeve, A. H., Kennedy, J. D., Marki, P. Z., Korneliussen, T. S., Freeman, B. G., Sam, K., Linck, E., Haryoko, T., Iova, B., Koane, B., Maiah, G., Paul, L., Irestedt, M., & Jønsson, K. A. (2022). The formation of avian montane diversity across barriers and along elevational gradients. Nature Communications, 13(1), 268.

68. Purcell, S., Neale, B., Todd-Brown, K., Thomas, L., Ferreira, M. A. R., Bender, D., Maller, J., Sklar, P., de Bakker, P. I. W., Daly, M. J., & Sham, P. C. (2007). PLINK: a tool set for whole-genome association and population-based linkage analyses. The American Journal of Human Genetics, 81(3), 559–575.

69. Qin, S.-Y., Zuo, Z.-Y., Xu, S.-X., Liu, J., Yang, F.-M., Luo, Y.-H., Ye, J.-W., Zhao, Y., Rong, J., Liu, B., Ma, P.-F., & Li, D.-Z. (2024). Anthropogenic disturbance driving population decline of a dominant tree in East Asia evergreen broadleaved forests over the last 11,000 years. Conservation Biology: The Journal of the Society for Conservation Biology, 38(1), e14180.

70. Quinlan, A. R., & Hall, I. M. (2010). BEDTools: a flexible suite of utilities for comparing genomic features. *Bioinformatics (Oxford*, England), 26(6), 841–842.

71. Quinn, C. B., Preckler-Quisquater, S., Buchalski, M. R., & Sacks, B. N. (2024). Whole genomes inform genetic rescue strategy for Montane red foxes in north America. Molecular Biology and Evolution, 41(9). 10.1093/molbev/msae193

72. Quintero, I., & Wiens, J. J. (2013). Rates of projected climate change dramatically exceed past rates of climatic niche evolution among vertebrate species. Ecology Letters, 16(8), 1095–1103.

73. Ramesh, V., Hariharan, P., Gupte, P. R., Mohan, A. V., Akshay, V. A., Rajan, A., Das, C., Lockwood, I., Robin, V. V., Tingley, M. W., & DeFries, R. (2025). Grassland bird species decline with colonial-era landscape change in a tropical Montane ecosystem. Global Change Biology, 31(7), e70358.

74. Ramirez, B. R., Kass, J. M., Zellmer, A. J., Hund, A. K., & Scordato, E. S. C. (2025). Long-term human land-use change throughout southeast Asia reshapes the distribution of suitable habitat for a human-commensal bird species. Journal of Biogeography, 52(6), Not Available.

75. Rice, C. G. (1984). The behavior and ecology of Nilgiri tahr (Hemitragus hylocrius Ogilby, 1838). https://oaktrust.library.tamu.edu/items/88540934-b54b-4104-95b4-176322b23039

76. Robin, V. V., & Nandini, R. (2012). Shola habitats on sky islands: status of research on montane forests and grasslands in southern India. Current Science, 103(12), 1427–1437.

77. Robin, V. V., Sinha, A., & Ramakrishnan, U. (2010). Ancient geographical gaps and paleo-climate shape the phylogeography of an endemic bird in the sky islands of southern India. PloS One, 5(10), e13321.

78. Robin, V. V., & Sukumar, R. (2002). Status and habitat preference of White-bellied Shortwing Brachypteryx major in the Western Ghats (Kerala and Tamilnadu), India. Bird Conservation International, 12(4), 335–351.

79. Robin, V. V., Sukumar, R., & Thiollay, J.-M. (2006). Status and distribution of the White-bellied Shortwing Brachypteryx major in the Western Ghats of Karnataka and Goa, India. Bird Conservation International, 16(4), 345–351.

80. Santiago, E., Köpke, C., & Caballero, A. (2025). Accounting for population structure and data quality in demographic inference with linkage disequilibrium methods. Nature Communications, 16(1), 6054.

81. Santiago, E., Novo, I., Pardiñas, A. F., Saura, M., Wang, J., & Caballero, A. (2020). Recent demographic history inferred by high-resolution analysis of linkage disequilibrium. Molecular Biology and Evolution, 37(12), 3642–3653.

82. Schiffels, S., & Durbin, R. (2014). Inferring human population size and separation history from multiple genome sequences. Nature Genetics, 46(8), 919–925.

83. Schiffels, S., & Wang, K. (2020). MSMC and MSMC2: The Multiple Sequentially Markovian Coalescent. Methods in Molecular Biology (Clifton, N.J.), Methods in Molecular Biology (Clifton, N.J.), 2090, 147–166.

84. Sharma, R., Arora, N., Goossens, B., Nater, A., Morf, N., Salmona, J., Bruford, M. W., Van Schaik, C. P., Krützen, M., & Chikhi, L. (2012). Effective population size dynamics and the demographic collapse of Bornean orang-utans. PloS One, 7(11), e49429.

85. Sirami, C., Nespoulous, A., Cheylan, J.-P., Marty, P., Hvenegaard, G. T., Geniez, P., Schatz, B., & Martin, J.-L. (2010). Long-term anthropogenic and ecological dynamics of a Mediterranean landscape: Impacts on multiple taxa. Landscape and Urban Planning, 96(4), 214–223.

86. Smeds, L., Qvarnström, A., & Ellegren, H. (2016). Direct estimate of the rate of germline mutation in a bird. Genome Research, 26(9), 1211–1218.

87. Sreekumar, E. R., & Nameer, P. O. (2022). A MaxEnt modelling approach to understand the climate change effects on the distributional range of White-bellied Sholakili Sholicola albiventris (Blanford, 1868) in the Western Ghats, India. Ecological Informatics, 70(101702), 101702.

88. Stapley, J., Feulner, P. G. D., Johnston, S. E., Santure, A. W., & Smadja, C. M. (2017). Variation in recombination frequency and distribution across eukaryotes: patterns and processes. Philosophical Transactions of the Royal Society of London. Series B, Biological Sciences, 372(1736). 10.1098/rstb.2016.0455

89. Stefanović, M., Bogdanowicz, W., Adavoudi, R., Martínez-Sosa, F., Doan, K., Flores-Manzanero, A., Srinivas, Y., Banea, O. C., Ćirović, D., D’Amico, G., Djan, M., Giannatos, G., Hatlauf, J., Hayrapetyan, V., Heltai, M., Homel, K., Hulva, P., Ionică, A. M., Jhala, Y. V., … Pilot, M. (2024). Range-wide phylogeography of the golden jackals (Canis aureus) reveals multiple sources of recent spatial expansion and admixture with dogs at the expansion front. Biological Conservation, 290(110448), 110448.

90. Stewart, J. R., Lister, A. M., Barnes, I., & Dalén, L. (2010). Refugia revisited: individualistic responses of species in space and time. *Proceedings*. Biological Sciences, 277(1682), 661–671.

91. Suresh, S., Rajesh, S., & Pradeepkumar, A. P. (2018). History and Development of Devikulam High Range, Idukki District, Kerala. Heritage: Journal of Multidisciplinary Studies in Archaeology, 6, 697–711.

92. Svenning, J.-C., Eiserhardt, W. L., Normand, S., Ordonez, A., & Sandel, B. (2015). The influence of paleoclimate on present-day patterns in biodiversity and ecosystems. Annual Review of Ecology, Evolution, and Systematics, 46(1), 551–572.

93. Teng, Y., Li, W., Wang, X., Zhang, R., Shen, Y., Wu, R., Liu, J., Zhang, M., Roos, C., Li, J., Li, J., Qi, J., & Li, M. (2025). Integrating population genomics and environmental data to predict adaptation to climate change in post-bottleneck Tibetan macaques. Science Advances, 11(28), eadw0562.

94. Thomas, C. D., Cameron, A., Green, R. E., Bakkenes, M., Beaumont, L. J., Collingham, Y. C., Erasmus, B. F. N., De Siqueira, M. F., Grainger, A., Hannah, L., Hughes, L., Huntley, B., Van Jaarsveld, A. S., Midgley, G. F., Miles, L., Ortega-Huerta, M. A., Peterson, A. T., Phillips, O. L., & Williams, S. E. (2004). Extinction risk from climate change. Nature, 427(6970), 145–148.

95. Thomas, N. E., Chadwick, E. A., Bruford, M. W., & Hailer, F. (2025). Spatio-temporal changes in effective population size in an expanding metapopulation of Eurasian otters. Evolutionary Applications, 18(1), e70067.

96. Tian, H., Banger, K., Bo, T., & Dadhwal, V. K. (2014). History of land use in India during 1880–2010: Large-scale land transformations reconstructed from satellite data and historical archives. Global and Planetary Change, 121, 78–88.

97. Tingley, M. W., Monahan, W. B., Beissinger, S. R., & Moritz, C. (2009). Birds track their Grinnellian niche through a century of climate change. Proceedings of the National Academy of Sciences of the United States of America, 106 Suppl 2(supplement_2), 19637–19643.

98. Vinay, K. L. (n.d.). BirdLab-Protocols: General procedures and protocols followed in BirdLab based at IISER Tirupati. Github. Retrieved March 25, 2026, from https://github.com/stachyris/BirdLab-Protocols?tab=readme-ov-file

99. Vinay, K. L., Arvind, C., Goyal, N., & Robin, V. V. (2025). High-quality genome assembly of the endemic, threatened, White-bellied Sholakili Sholicola albiventris (Muscicapidae: Blanford, 1868) from the Shola Sky Islands, India. The Journal of Heredity, (esaf049). 10.1093/jhered/esaf049

100. Viswanathan, A., Thrikkadeeri, K., Koulgi, P., Praveen, J., Deomurari, A., Jha, A., Warudkar, A., Suryawanshi, K., Madhusudan, M. D., Kaushik, M., Goyal, N., Bangal, P., Jayapal, R., Quader, S., Dutta, S., Menon, T., & Ramachandran, V. (2024). State of India’s Birds 2023: A framework to leverage semi-structured citizen science for bird conservation. In bioRxiv. bioRxiv. 10.1101/2024.09.05.611348

101. von Seth, J., van der Valk, T., Lord, E., Sigeman, H., Olsen, R.-A., Knapp, M., Kardailsky, O., Robertson, F., Hale, M., Houston, D., Kennedy, E., Dalén, L., Norén, K., Massaro, M., Robertson, B. C., & Dussex, N. (2022). Genomic trajectories of a near-extinction event in the Chatham Island black robin. BMC Genomics, 23(1), 747.

102. Wang, J., Santiago, E., & Caballero, A. (2016). Prediction and estimation of effective population size. Heredity, 117(4), 193–206.

103. Warudkar, A. (2025). Emerging infectious disease in the Shola sky islands. Indian Institute of Science Education and Research Tirupati.

104. Wright, S. (1931). Evolution in Mendelian populations. Genetics, 16(2), 97–159.

