## Supplementary methods & results for "Recovering the Impacts of Landscape Changes through the Demographic History of a Western Ghats Endemic Bird"

### S2.2.4 Genomic Data Preprocessing

The adapter sequences and low-quality reads were then trimmed out using *Trimmomatic v0.39* (Bolger et al., 2014) in paired-end mode with -phred33 encoding. Illumina TruSeq3 adapters (ILLUMINACLIP:TruSeq3-PE.fa:2:30:10:2:True), low-quality bases from leading (LEADING:3) and trailing (TRAILING:3) ends, and reads shorter than 36 bp (MINLEN:36) were removed during trimming. The raw and trimmed reads were inspected for sequence quality, GC content, and adapter contamination using *FastQC v0.12.1* (*Babraham Bioinformatics - FastQC A Quality Control Tool for High Throughput Sequence Data*, n.d.). The scaffold-level reference assembly for White-bellied Sholakili was downloaded from NCBI genomes (GCA_050043525.1; Scaffold N50: 68.6 Mb) (Vinay et al., 2025). The reference was indexed using *bwa v0.7.18-r1243 (Li, 2013)* with the index option. The trimmed paired reads were then mapped to the WBS reference assembly using *bwa v0.7.18-r1243* with the mem option (Li, 2013), marking shorter split hits as secondary (-M), and adding Read group information, including read group ID, Sample name, library, and platform (-R "@RG\tID:ID\tSM:name\tLB:IlluminaWGS\tPL:ILLUMINA"). The SAM output was then converted to BAM format and sorted using *samtools v1.20* (Li et al., 2009). The mapped raw BAMs were then filtered for unmapped reads (-F 4) and low-quality mapping (-q 30) using *samtools*. The filtered BAMs were then resorted using *samtools*. The PCR duplicates were identified and removed using *Picard Tools v2.27.5 MarkDuplicates* (*Picard*, 2022), with metrics recorded for the samples. The final BAMs were indexed using *samtools* with the index option. The quality of the BAMs was assessed using *Qualimap v2.3 bamqc* (Okonechnikov et al., 2016) to evaluate genomic coverage and mapping statistics for each of the raw, filtered, and PCR-duplicate-removed BAMs.

### S2.2.5 Variant calling and filtering:

Further analyses were conducted at the population level, repeating the downstream steps for each of the four populations. The variant sites from the final BAMs were identified independently using both *bcftools v1.10.2* (Li, 2012) and *freebayes v1.3.6* (Garrison & Marth, 2012). In the bcftools pipeline, the genotype likelihoods were identified first from the BAM files using the *bcftools mpileup* function with annotations for allelic depth (AD), number of high-quality bases (read depth) (DP), and phred-scaled strand bias p-value (SP). SNPs and Indels were called using the bcftools call function with the multiallelic calling model (-m), retaining genotype likelihoods (GQ) and posterior probabilities (GP). In the freebayes pipeline, the variant sites were identified using the same BAMs and reference with default parameters. To obtain only the high-confidence sites, the variant sites recognised by both bcftools and freebayes were retained using *bedtools v2.31.1 intersect* (Quinlan & Hall, 2010). The intersected VCF was compressed and indexed for downstream processing.

The intersected VCF was then filtered to retain only biallelic SNPs using *bcftools v2.31.1* (Danecek et al., 2021), and indels were removed using *vcftools v0.1.16* (Danecek et al., 2011). Additional stringent filtering was done using *vcftools* to retain only the high-quality variant sites. The filters used were: minimum genotype quality (--minGQ 30), minimum and maximum read depth (--minDP 5 --maxDP 30), minor allele frequency (--maf 0.05), Hardy-Weinberg equilibrium (--hwe 0.05), and maximum missingness (--max-missing 0.9).

The small scaffolds (< 20 Mb) and the sex linked scaffolds (scaffold33 and scaffold35 - identified using the approach in Natesh et al. (in prep. )) were removed from the filtered VCF, retaining only the big scaffolds (scaffolds1 - scaffold14). The final filtered VCFs were converted to PED and MAP formats using *PLINK v1.90b7.2* (Purcell et al., 2007).

For analyses other than demographic estimations, regions in Linkage Disequilibrium (LD) were identified with the *PLINK* sliding-window approach (--indep-pairwise 50 10 0.1) and pruned with *vcftools*.

### S2.3.3 Estimating the recombination rate of White-bellied Sholakili

Recombination rates required for demographic inferences in *GONE2* were estimated with *iSMC* (Barroso et al., 2019). For the analysis, the individual with the highest average sequencing depth was selected from the KD population. The analysis is restricted to include only scaffolds larger than 100 kb, as shorter scaffolds provide limited coalescent-based information. The *bamCaller.py* scripts from *msmc-tools* pipeline (Schiffels & Wang, 2020) were used to do variant calling and mask generation. Variants were called independently for each scaffold using *bcftools mpileup* and *call* functions with stringent filters, including probabilistic realignment for base alignment quality (BAQ) disabled (-B), minimum mapping quality (-q 20), minimum base quality (-Q 20), adjustment for mapping quality (-C 50), and skipping indels (-V indels). Masks were generated using *bamCaller.py*, applying a depth filter based on the calculated mean depth. The VCFs and corresponding mask files identifying high-confidence callable regions were generated as outputs.

The VCF files generated for each scaffold were concatenated and sorted. A tab-separated file specifying scaffold IDs and the start and end coordinates was generated using the *generate_tab.py* scripts from the *iSMC* pipeline. The mask files from all scaffolds were merged and sorted to generate a genome-wide mask file of all the callable regions. A “negative mask” complementary to the generated mask was prepared by subtracting the callable regions from the full genome coordinates using bedtools subtract. This mask was further trimmed to include only the regions present in the tab file.

An options file for *iSMC* was created, using the prepared genome-wide VCF, tab file, and negative mask file as inputs and the following parameters: diploid_indices = (0,1), init_number_knots = 2, init_number_knots = 3, number_rho_categories = 5, number_intervals = 40, function_tolerance = 1e-4, and fragment_size = 3000000. iSMC was run with parameter optimisation, posterior decoding, and sequence summarising. The output rho (population-scaled recombination rate; ρ) estimates were averaged in windows of 10kb, 100kb, and 1 Mb using the ismc_mapper function. The Ne value for the KD population was calculated from Watterson’s theta ($\Theta_w$), estimated using a modified version of *genomics_vcf_summary_stats.py* from (Funk,

2024). The rho value was converted to recombination rate (r) in cM/Mb using the following formula.

$\rho = 4*Ne*r$ ; r = recombination rate in crossovers per base pair per generation
$r\_cMMb = r*10e8$ ; r_cMMb = recombination rate in cM/Mb

$\Theta_w = 4*Ne*\mu$ ; μ = mutation rate

### S2.4 Validating demographic history estimations using genomic simulations

Information from the demographic history patterns of four populations (HW, BJ, KD, and PV - selected based on differing patterns of demography) was used as input for *SLiM v5.0* (Haller et al., 2025) to run genomic simulations. All the simulation runs were initialised with a nucleotide-based model (initializeSLiMOptions(nucleotideBased=T)) and a multi-chromosome model (initializeChromosome()) using 14 chromosomes, each of a similar size to the first 14 scaffolds of the White-bellied Sholakili reference. The simulations utilised only genomic elements with neutral nucleotide-based mutations (dominance coefficient = 0.5; selection coefficient with fixed distribution = 0), with a species-specific mutation rate of 7.31e-9, implemented via a Jukes-Cantor model matrix (mmJukesCantor(7.31e-9)). The species-specific mutation rate for the same was estimated from the generation time (3.18 years) of Bird et al. (2020) and the per-site-per-year mutation rate of Collard Flycatchers (CF) (2.3e-9) from Smeds et al. (2016) using the equation '*mutation rate per site per generation in WBS = generation time of WBS * mutation rate per site per year in CF (approximately)*'. The default recombination rate of 1e-8 crossovers per base pair per generation (1 cM/Mb) was used for the simulations. Each simulation was run for an initial burn-in of 10000 ticks (generations) followed by another 1000 ticks, during which tweaks to subpopulation sizes and numbers were applied as needed. At the end of the last tick (11000th tick), chromosome-level VCFs of randomly selected 20 individuals from the final population (or [*20/number of population*] individuals from each final population) were generated as output using the outputHaplosomesToVCF function. These VCFs were combined into a single VCF using *bcftools concat*, filtered to remove invariant sites, and indexed. The final VCF was converted to PED and MAP formats using *PLINK* and then used to run *GONE2* with the same parameters as for demographic analysis. The results were visualised alongside the real data and simulation input to compare and validate the observations from real populations.

For each population, two sets of simulations were run: one with a real increase or decrease in Ne (hereafter, the 'baseline model'), and the other with population fusion or fragmentation, to assess potential artefacts in the dataset.

To validate the recent Ne increase pattern in the KD and BJ populations, two sets of simulations were run: first, the baseline model with an actual population increase and second, a population fusion model. For the baseline model, a single subpopulation in SLiM was simulated, varying its population sizes from approximately the 10950th generation to match one of the KD or BJ bootstraps. For the population fusion model, 2 and 4 subpopulations were simulated in the KD and BJ-based models, respectively, from the 10000th generation onwards. The population size was varied to match the true population until their point of increase, at which all subpopulations were combined into a single subpopulation. Results from initial runs were used to make tweaks and rerun the SLiM models until a satisfactory level of similarity to the real population was achieved.

To validate the recent Ne decline pattern in HW and KD populations, two sets of simulations were run: first, the baseline model with an actual population decline and second, a population fragmentation model. For the baseline model, a similar approach to that used for the population increase model was applied. For the population fragmentation model, a single population was simulated to undergo changes similar to those in real populations until its decline, at which point a subpopulation split was simulated, forming two subpopulations. Similar to earlier, results from initial runs were used to make tweaks and rerun the SLiM models until a satisfactory level of similarity to the real population was achieved.

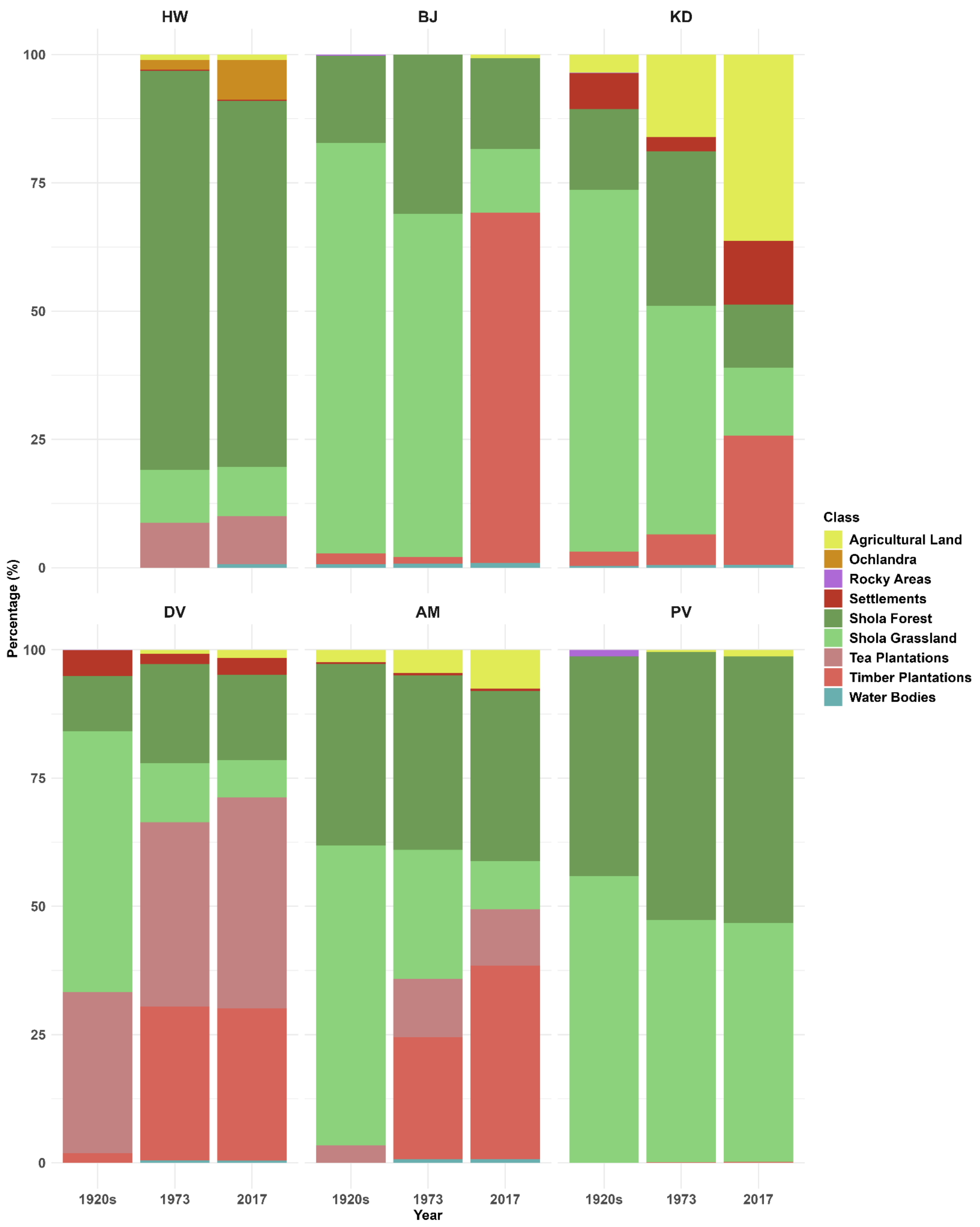

*Figure S1a: Stacked bar plot of the percentage of each land cover class in a 5km radius buffer around the sampling sites across 6 populations over 3 time points. Data unavailable for HW from the 1920s. Classification of the Rocky area is absent for 1973 & 2017. Made from the historical maps of Lockwood et. al. (in prep); Arasumani et al. (2019).*

A)

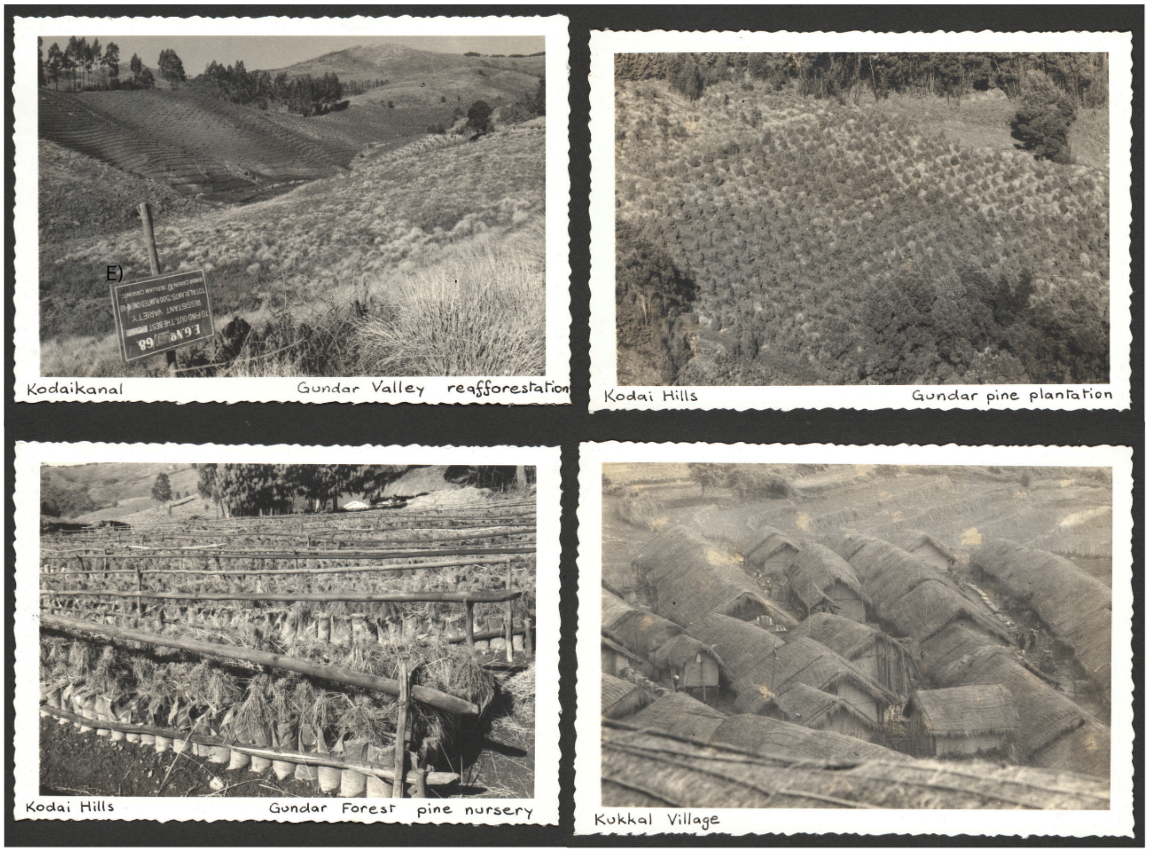

B)

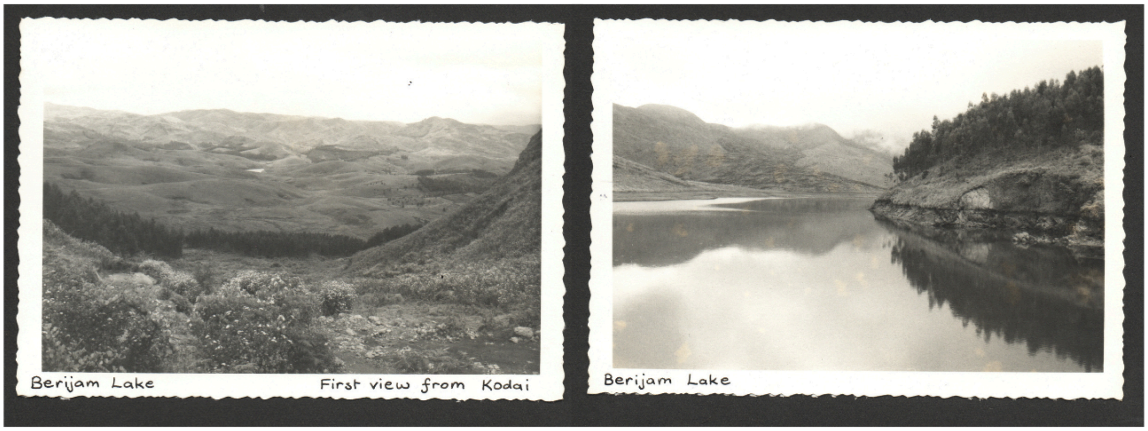

*Figure S1b: Archived historic photographs of Kodaikanal (A) and Berijam (B) from the 1960s, indicating the presence of timber plantations, villages, and agriculture in Kodaikanal and Timber plantations in Berijam. Images taken by Dr Nora Mitchell; Images acquired from Ian Lockwood.*

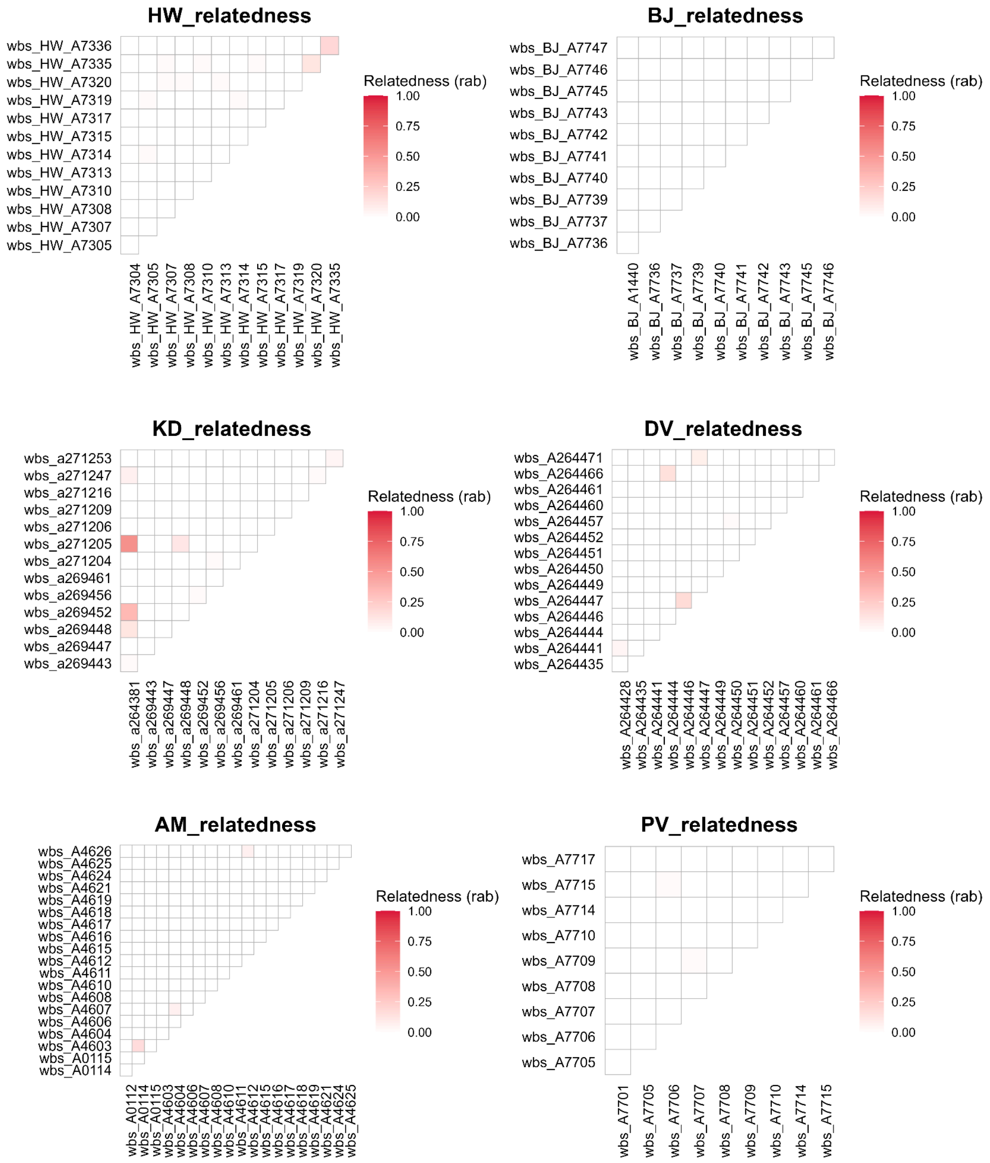

*Figure S2: Relatedness heatmaps: HW - HighWavies; BJ - Berijam; KD - Kodaikanal; DV - Devikulam; AM - Anamudi; PV - Poovar. The colour scale represents Pairwise Relatedness (rab) from the ngsRelate output, with darker colours implying greater relatedness.*

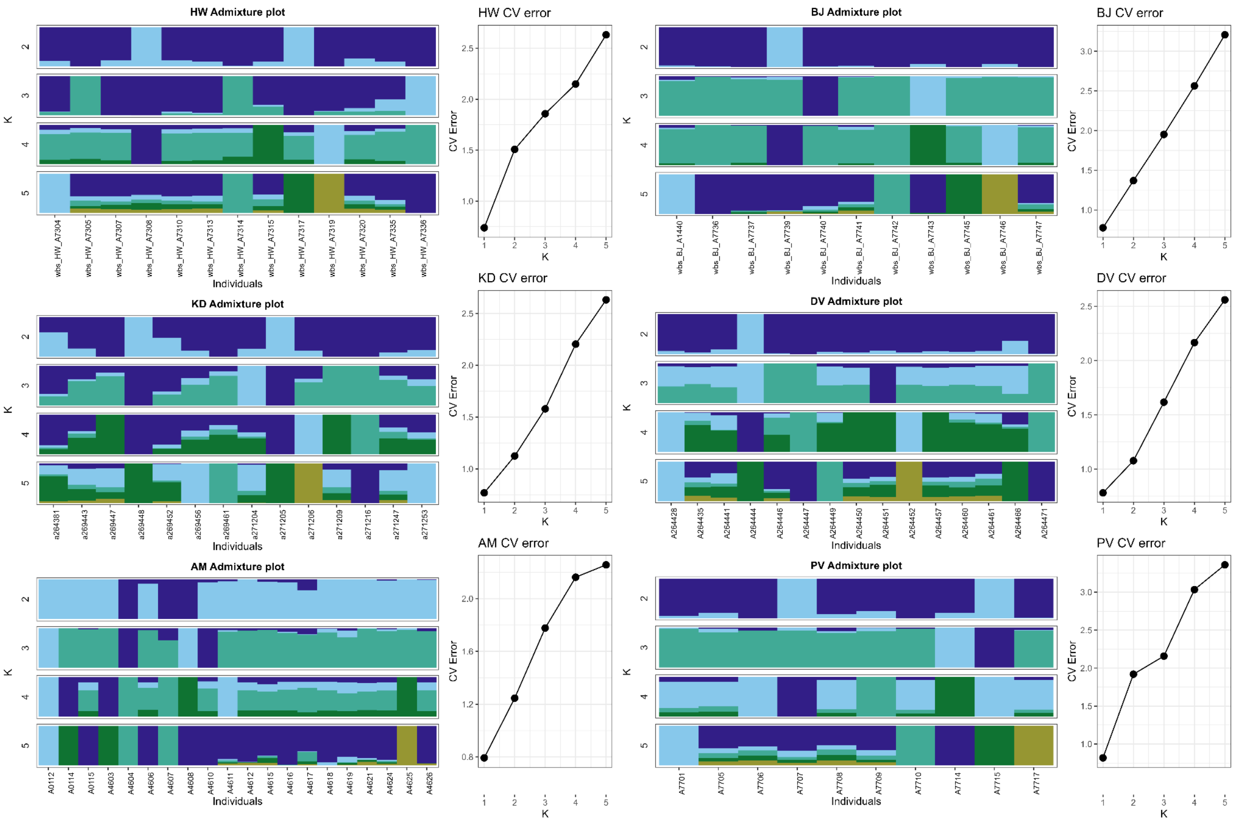

*Figure S3: ADMIXTURE outputs for each population. Admixture barplots showing proportion of ancestry for the Number of clusters, K=2-5. The vertical bars represent individuals, and each colour corresponds to a genetic cluster inferred by the software. Cross-validation (CV) error plots for K values from 1 to 5 for each population are attached to the admixture plots. The smallest CV error corresponds to the K value with the best support.*

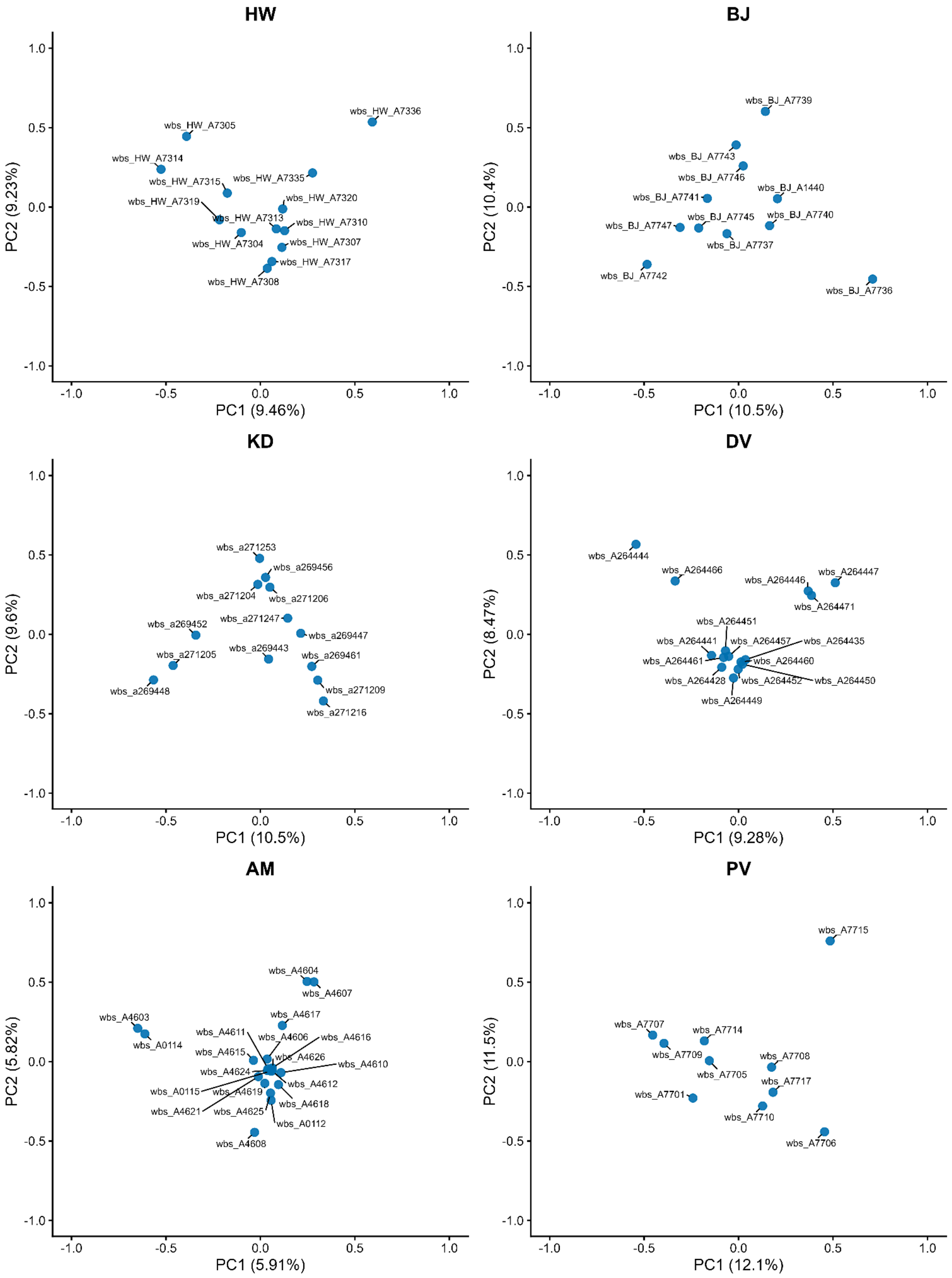

*Figure S4: PCA results for each population. The PC1 and PC2 are plotted. The weightage of each principal component is represented in parentheses beside the PCs. Each dot represents a single individual, labelled next to it.*

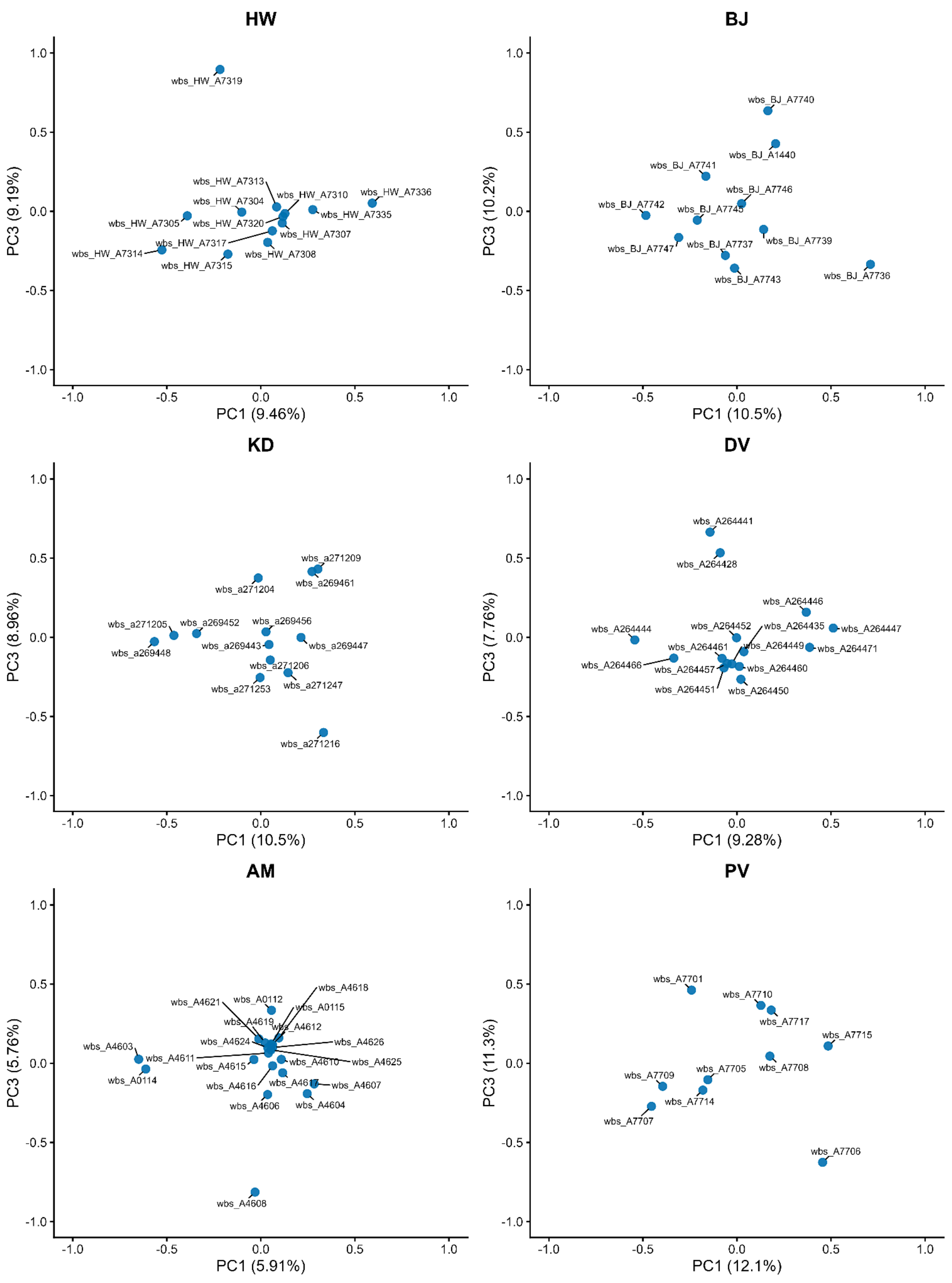

*Figure S5: PCA results for each population across PC1 and PC3. The weightage of each principal component is represented in parentheses beside the PCs. Each dot represents a single individual, labelled next to it.*

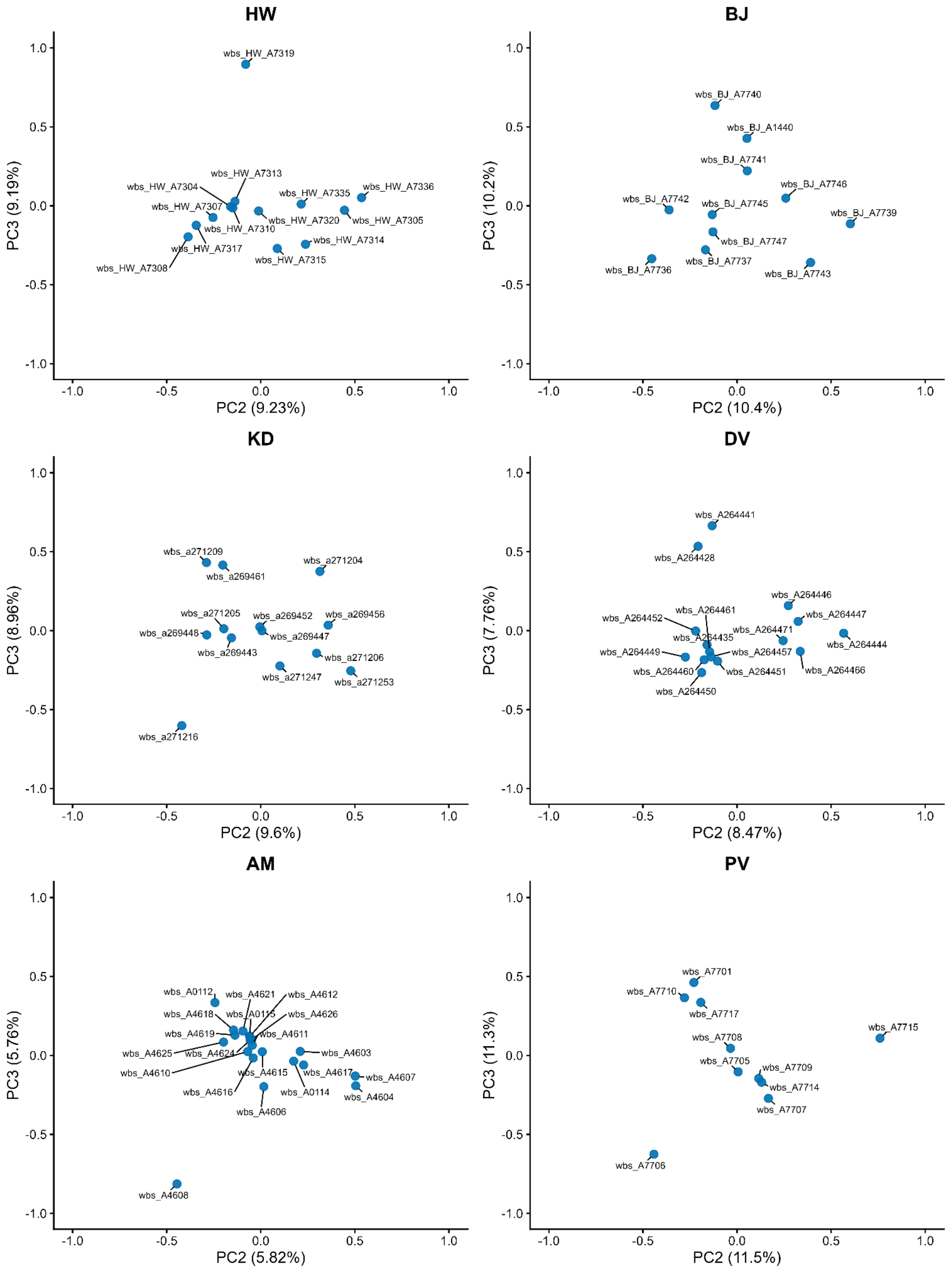

*Figure S6: PCA results for each population across PC2 and PC3. The weightage of each principal component is represented in parentheses beside the PCs. Each dot represents a single individual, labelled next to it.*

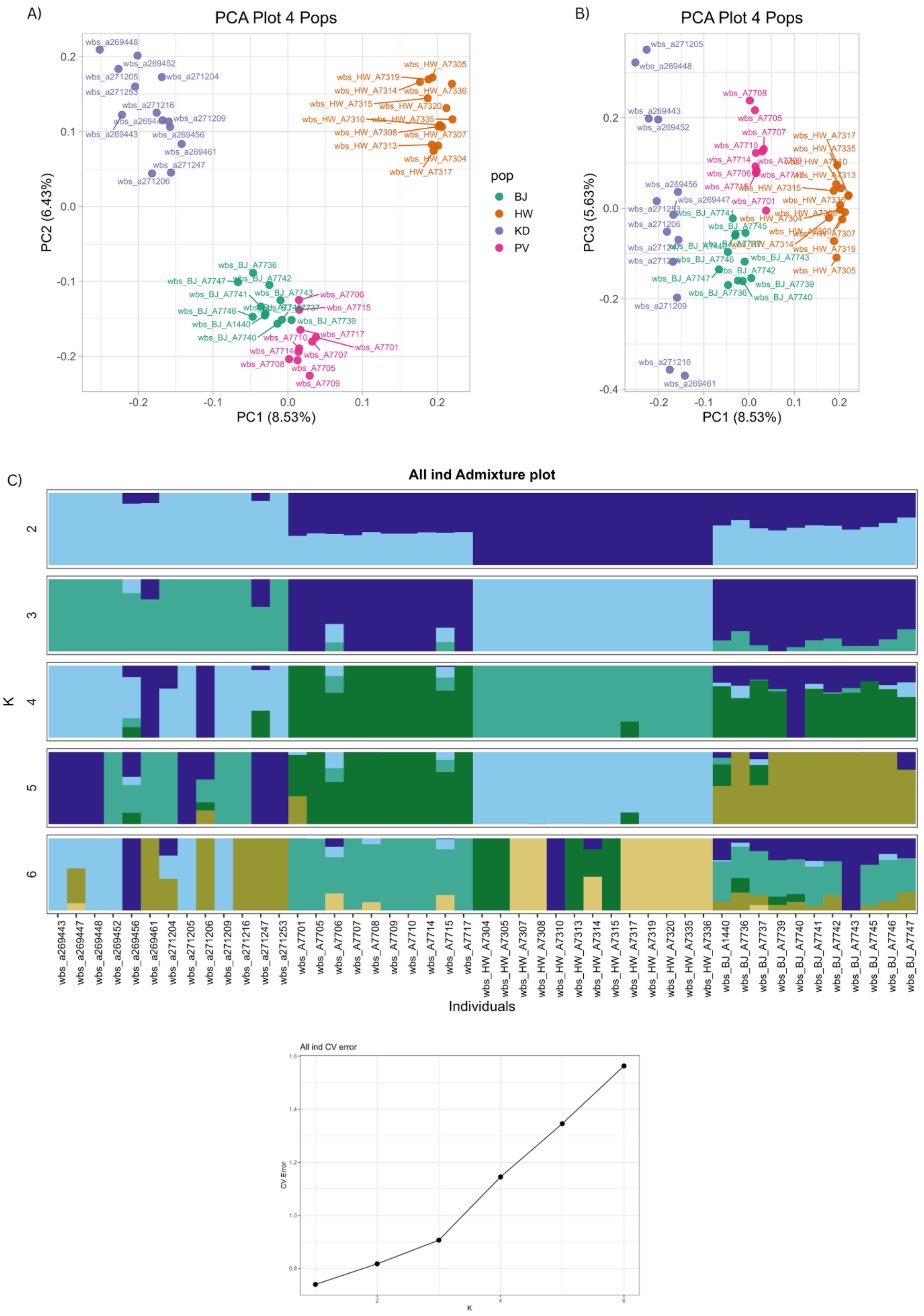

*Figure S7: PCA and ADMIXTURE plots for four populations (HW, BJ, KD, PV)*

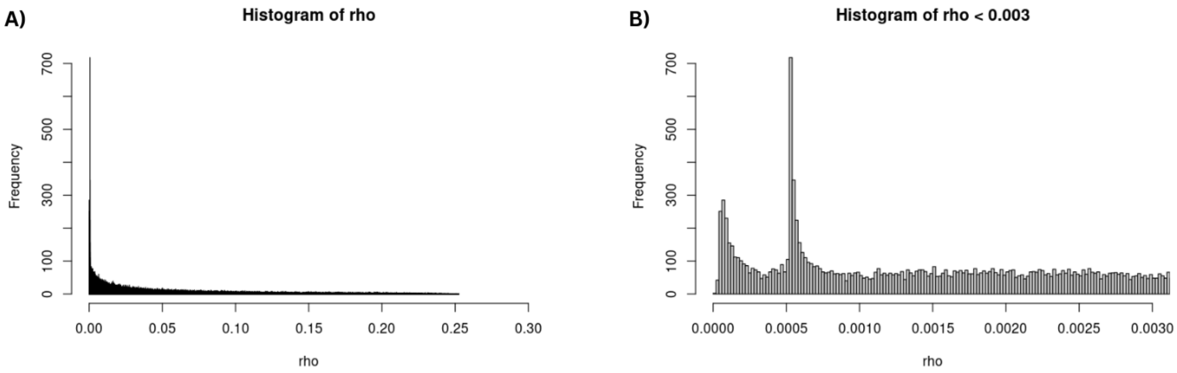

**Mean = 0.045**
**Median = 0.023**

**Most Frequent = 0.00053**

*Figure S8: The rho estimates averaged to bins of 10kb from iSMC. A) Histogram of all the rho values estimated by iSMC across the whole genome - 1000 bins. Mean and Median values of the distribution given below the figure. B) Histogram of rho values less than 0.003. The most frequent value is given below the figure.*

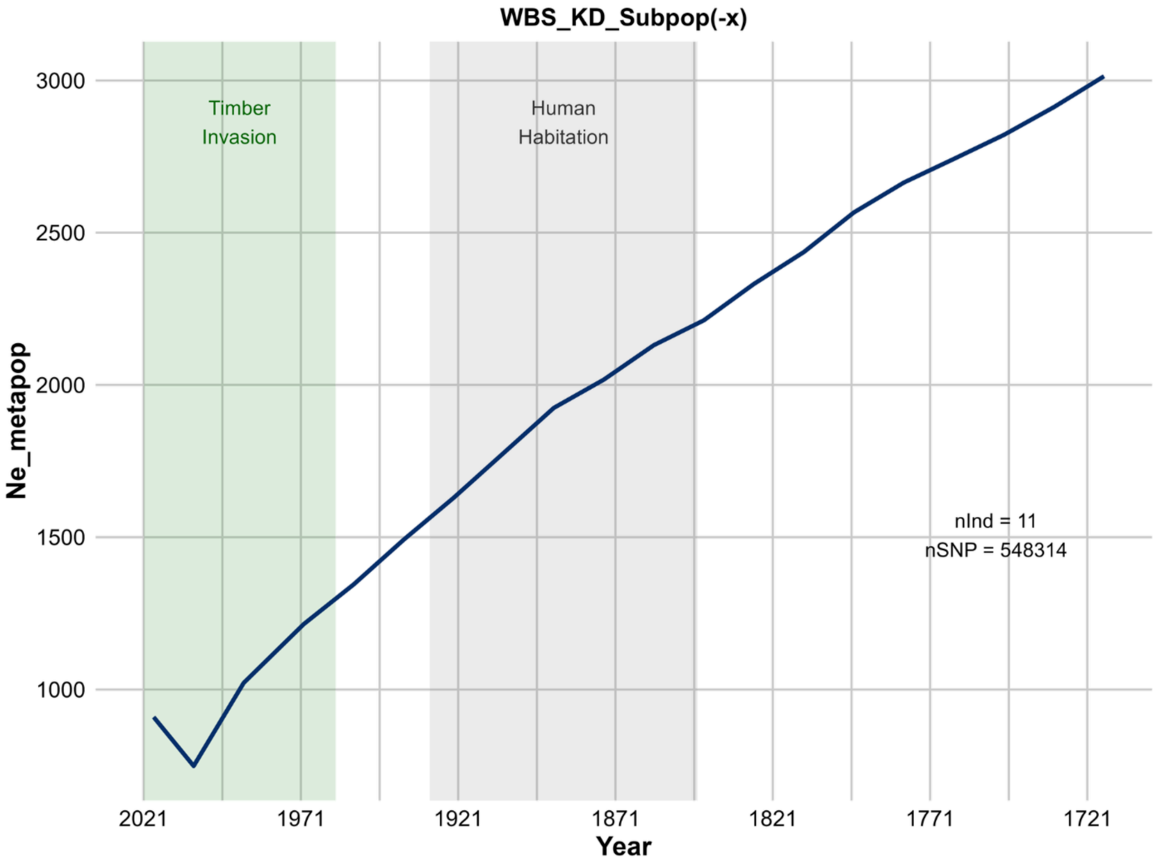

*Figure S9: The GONE2 results on the KD population with a subpopulation assumption (-x). X-axis: Ne_metapop - Ne of the entire metapopulation containing two subpopulations.*

HW Simulations

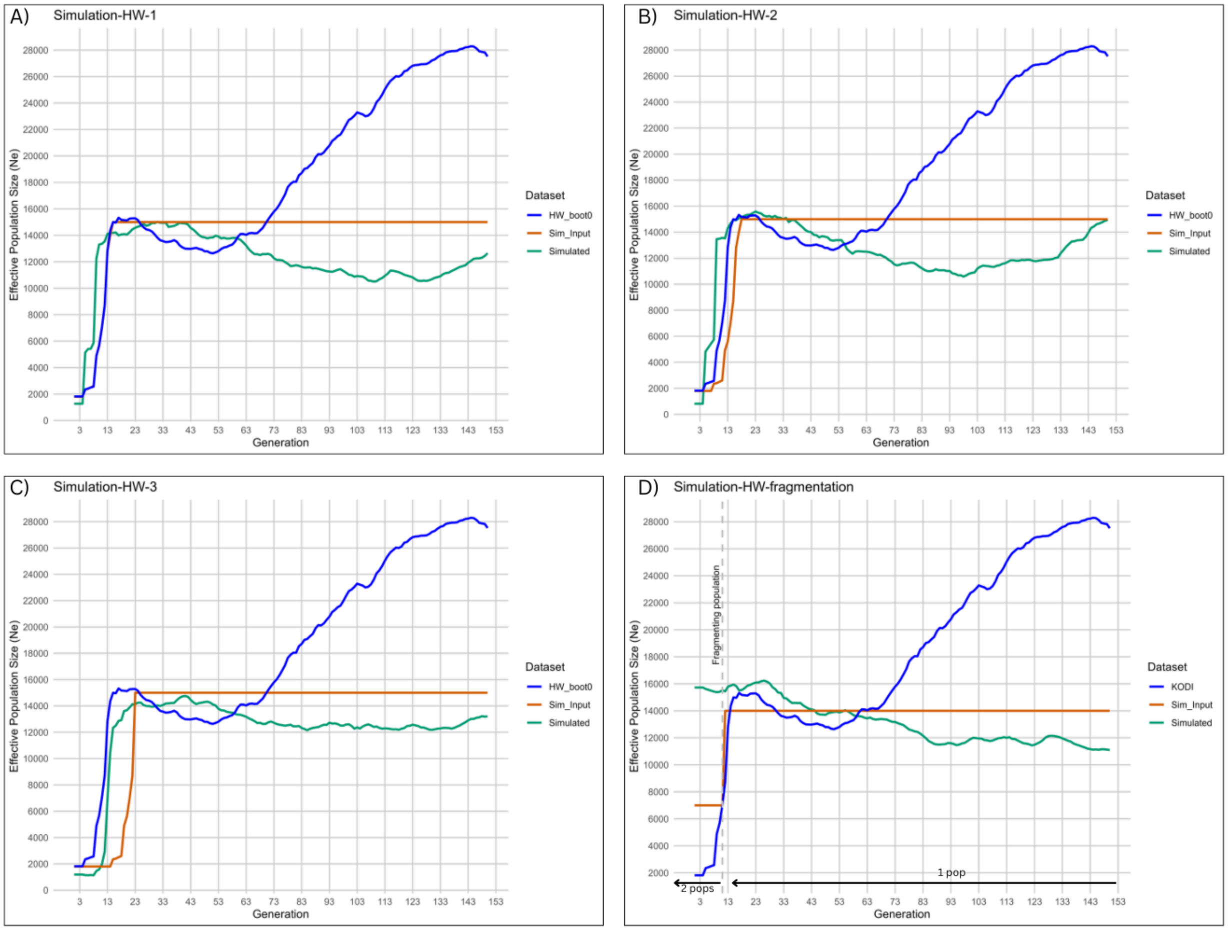

*Figure S10: SLiM simulations on HW population. The dark blue line is the first GONE2 bootstraps (boot0) from the HW population. The orange line is the input given in SLiMgui. In simulations with the fragmentation model, the orange line represents the input population size of a single subpopulation. Green is the estimated Ne output of GONE2 from 20 individuals randomly selected from 'Generation 0' of the simulated dataset. A) Baseline model: simulations with the inputs similar to the HW bootstrap. B-C) Simulations with Ne change, similar to the HW bootstrap, but with the timing of population decline varied. D) Simulations with one subpopulation getting fragmented into two. The number and timeline of subpopulations are represented by black arrows.*

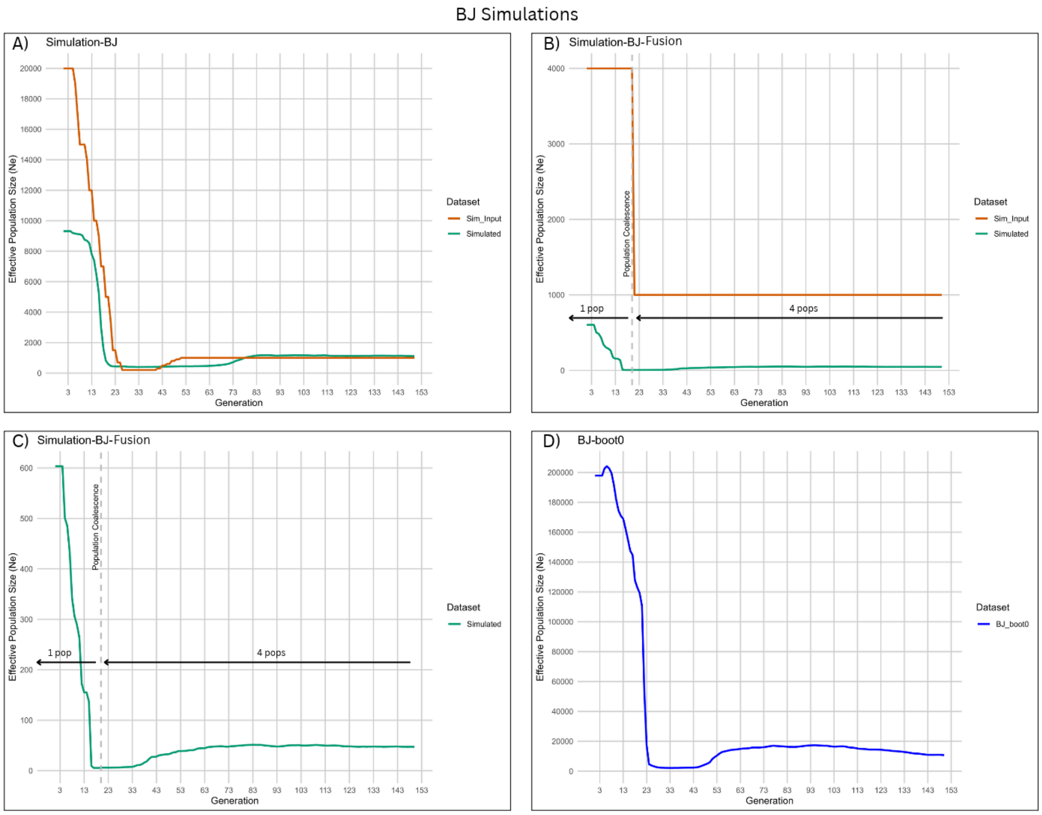

*Figure S11: SLiM simulations on the BJ population. The dark blue line is the first GONE2 bootstraps (boot0) from the BJ population. The orange line is the input given in SLiMgui. In simulations with the fusion model, the orange line represents the input population size of a single subpopulation. Green is the estimated Ne output of GONE2 from 20 individuals randomly selected from 'Generation 0' of the simulated dataset. A) Baseline model: simulations with the fold change of Ne increase similar to the BJ population. B) Simulations with four subpopulations getting coalesced into one. The number and timeline of subpopulations are represented by black arrows. C)The output from B without the simulation input. D) Bootstrap 0 from the BJ population.*

KD Simulations

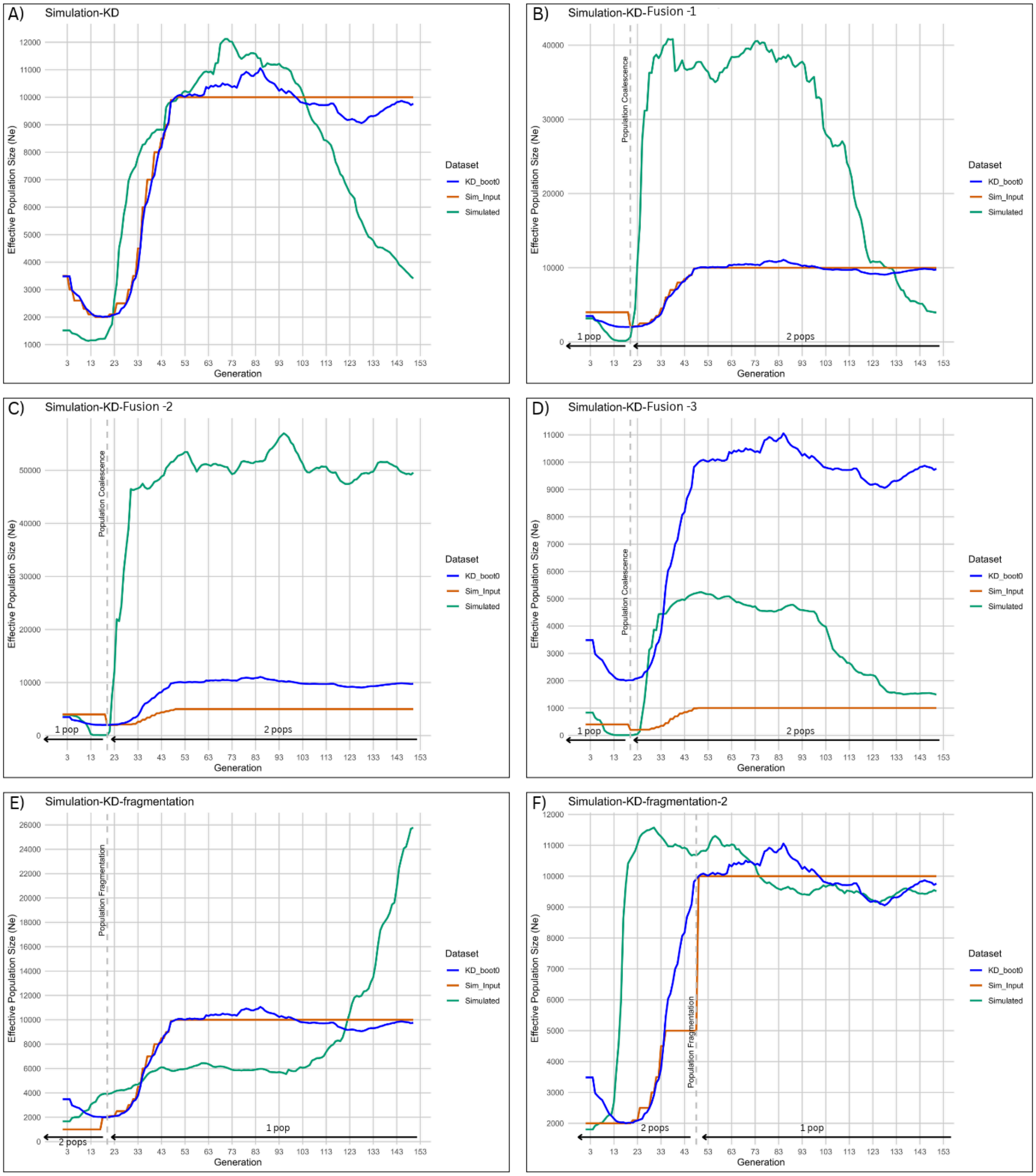

*Figure S12: SLiM simulations on KD population. The dark blue line is the first GONE2 bootstraps (boot0) from the KD population. The orange line is the input given in SLiMgui. In fusion and fragmentation model simulations, the orange line is the input population size of a single subpopulation. The green line is the output of GONE2 from 20 individuals randomly selected from 'Generation 0' of the simulated dataset. A) Baseline model: simulations with the inputs similar to the KD bootstrap. B-D) Simulations with two subpopulations getting coalesced*

*into one. The number and timeline of subpopulations are represented by black arrows. E-F) Simulations with one subpopulation getting fragmented into two. The number and timeline of subpopulations are represented by black arrows.*

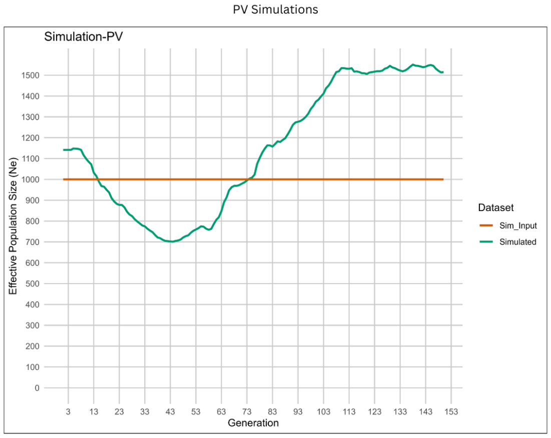

*Figure S13: SLiM simulations on the PV population - Baseline model. The orange line is the input given in SLiMgui. Green is the estimated Ne output of GONE2 from 20 individuals randomly selected from 'Generation 0' of the simulated dataset.*

*Table S1: FastQC and BamQC outputs for each sample. Gbp: Gigabasepairs; SD: Standard Deviation; X: fold coverage. The sample coloured in red is excluded from the downstream analyses. Samples coloured in xxx were excluded from the demographic analyses to maintain pamixia.*

| **Population** | **Ring No.** | **Raw (Gbp)** | **Trimmed (Gbp)** | **Mapped bam** | | **Filtered bam** | | **PCR duplicate removed bam** | |
|---|---|---|---|---|---|---|---|---|---|
| | | | | **Mean Coverage (X)** | **SD Coverage (X)** | **Mean Coverage (X)** | **SD Coverage (X)** | **Mean Coverage (X)** | **SD Coverage (X)** |
| HW | A7304 | 21.4 | 21 | 19.4426 | 120.7734 | 17.9145 | 41.7069 | 17.0312 | 21.1199 |
| | A7305 | 16 | 15.8 | 14.6333 | 119.6 | 13.2025 | 39.8874 | 12.7371 | 18.8682 |
| | A7307 | 17.4 | 17.2 | 15.836 | 120.4985 | 14.3886 | 35.3507 | 13.7676 | 19.3294 |
| | A7308 | 20.6 | 20.4 | 18.8608 | 103.2935 | 17.4796 | 32.4643 | 16.8652 | 19.8028 |
| | A7310 | 22.2 | 22 | 20.3271 | 153.4766 | 18.4847 | 49.7281 | 18.0678 | 25.0387 |
| | A7313 | 24.6 | 24.2 | 22.3532 | 167.0469 | 20.3302 | 62.2887 | 19.8381 | 28.314 |
| | A7314 | 20.4 | 20.2 | 18.6403 | 110.6533 | 17.1419 | 31.2209 | 16.2745 | 19.0736 |

| | | | | | | | | | |
|---|---|---|---|---|---|---|---|---|---|
| | A7315 | 22.4 | 22 | 20.3803 | 124.8836 | 18.7993 | 42.4195 | 18.3473 | 23.1648 |
| | A7317 | 26.4 | 26.2 | 24.167 | 134.0939 | 22.3582 | 44.1882 | 21.9661 | 26.7705 |
| | A7319 | 19 | 18.8 | 17.4496 | 72.1195 | 16.3158 | 25.4766 | 15.8685 | 17.2249 |
| | A7320 | 27.6 | 27.4 | 25.3053 | 168.5321 | 23.1005 | 52.6067 | 22.6077 | 28.3111 |
| | A7335 | 24 | 23.8 | 21.994 | 158.4942 | 20.0754 | 49.3751 | 19.7072 | 27.8152 |
| | A7336 | 19 | 18.8 | 17.3949 | 155.114 | 15.6084 | 53.8999 | 15.2695 | 25.2495 |
| BJ | A1440 | 25 | 24.8 | 22.9123 | 296.0468 | 19.8224 | 96.5799 | 18.9511 | 37.1259 |
| | A7736 | 18.4 | 18 | 16.5703 | 217.8451 | 14.3007 | 83.3663 | 13.6163 | 28.0268 |
| | A7737 | 22 | 21.8 | 20.1008 | 270.2391 | 17.3815 | 90.0923 | 16.5549 | 32.2417 |
| | A7739 | 17 | 16.8 | 15.5935 | 177.895 | 13.6592 | 49.6509 | 13.0726 | 24.8149 |
| | A7740 | 19.4 | 19.2 | 17.7795 | 128.2221 | 16.1564 | 37.025 | 14.795 | 20.0618 |
| | A7741 | 24.8 | 24.6 | 22.7517 | 199.1523 | 20.3671 | 57.6356 | 19.6049 | 30.3089 |
| | A7742 | 16.8 | 16.8 | 15.4381 | 100.4599 | 14.1032 | 30.7791 | 13.644 | 19.5789 |
| | A7743 | 18 | 18 | 16.6367 | 112.8219 | 15.18 | 29.9685 | 14.5942 | 19.5975 |
| | A7745 | 20.2 | 20 | 18.4887 | 223.215 | 16.031 | 61.4538 | 15.4852 | 30.146 |
| | A7746 | 24.8 | 24.6 | 22.6752 | 300.5562 | 19.5951 | 95.0169 | 18.7567 | 37.0283 |
| | A7747 | 21.8 | 21.4 | 19.8111 | 271.7245 | 17.0586 | 101.5806 | 16.2746 | 33.5722 |
| KD | A264381 | 33.2 | 30.1 | 28.5457 | 221.3646 | 25.8039 | 86.3493 | 24.6874 | 31.1279 |
| | A269443 | 23.6 | 22 | 20.313 | 86.1647 | 18.9141 | 36.9583 | 18.2621 | 21.772 |
| | A269447 | 15.8 | 14.8 | 13.6717 | 75.445 | 12.6383 | 30.7516 | 12.2567 | 17.4556 |
| | A269448 | 16.6 | 14.4 | 13.2304 | 45.8087 | 12.4097 | 22.2909 | 11.9748 | 14.052 |
| | A269452 | 21.8 | 20.4 | 18.8965 | 177.3026 | 16.8187 | 70.7532 | 16.2789 | 28.2327 |
| | A269456 | 22 | 20.8 | 19.0917 | 110.6177 | 17.4547 | 43.5294 | 16.9163 | 23.149 |
| | A269457 | 8.4 | 7.4 | 6.8576 | 37.5946 | 6.3247 | 15.1998 | 5.0746 | 7.9602 |
| | A269461 | 26.6 | 25 | 23.0557 | 169.0082 | 20.7987 | 67.8503 | 20.093 | 29.6628 |
| | A271204 | 15.2 | 14.4 | 13.3095 | 57.2024 | 12.4011 | 21.9897 | 11.9717 | 13.8861 |
| | A271205 | 32 | 30 | 27.5381 | 175.3629 | 25.2782 | 66.5195 | 24.1017 | 27.6653 |
| | A271206 | 14.8 | 13.2 | 12.1966 | 136.0705 | 10.6674 | 57.4667 | 10.2274 | 21.4507 |
| | A271209 | 19.8 | 18 | 16.6347 | 197.1597 | 14.436 | 71.0491 | 10.4828 | 19.6385 |
| | A271216 | 18.4 | 16 | 14.8366 | 207.4058 | 12.7534 | 80.8488 | 11.6454 | 21.7963 |
| | A271247 | 18 | 16.6 | 15.2692 | 247.9484 | 12.7357 | 94.7206 | 12.0326 | 29.7518 |
| | A271253 | 18.2 | 16.6 | 15.3265 | 47.9018 | 14.5056 | 21.6186 | 13.5511 | 14.2575 |

| | | | | | | | | | |
|---|---|---|---|---|---|---|---|---|---|
| DV | A264428 | 17.6 | 16.4 | 15.0503 | 95.3432 | 13.8094 | 45.4666 | 13.3971 | 22.3608 |
| | A264435 | 19.4 | 18.8 | 17.3366 | 125.2523 | 15.8406 | 49.0606 | 15.3474 | 26.4762 |
| | A264441 | 19.2 | 18.6 | 16.4954 | 155.2153 | 14.6124 | 57.4179 | 14.1712 | 26.8094 |
| | A264444 | 16.8 | 16 | 13.2402 | 107.8391 | 11.6702 | 38.0852 | 11.3807 | 20.0422 |
| | A264446 | 19.6 | 19 | 16.8315 | 141.6863 | 15.0493 | 52.0447 | 14.6121 | 25.2184 |
| | A264447 | 18.4 | 17.6 | 15.7337 | 140.4172 | 14.0233 | 53.2792 | 13.6535 | 25.8202 |
| | A264449 | 19 | 18.4 | 15.8131 | 134.1476 | 14.1195 | 47.7175 | 13.7372 | 23.6363 |
| | A264450 | 15.4 | 15 | 13.8313 | 171.7241 | 12.0456 | 67.9523 | 11.689 | 31.9473 |
| | A264451 | 15.8 | 15.2 | 14.0754 | 112.8253 | 12.6639 | 43.6225 | 12.2504 | 24.2262 |
| | A264452 | 18 | 17.4 | 15.7518 | 111.4867 | 14.3332 | 43.3527 | 13.9702 | 23.3812 |
| | A264457 | 16.8 | 16.2 | 15.0377 | 65.8754 | 14.124 | 28.1199 | 13.7954 | 18.8555 |
| | A264460 | 17.4 | 16.8 | 15.4857 | 159.0442 | 13.7539 | 63.9192 | 13.3091 | 30.3857 |
| | A264461 | 19 | 18.4 | 16.9633 | 92.3758 | 15.6748 | 36.4289 | 15.1942 | 22.5293 |
| | A264466 | 18 | 17.4 | 16.078 | 108.4214 | 14.7461 | 44.4321 | 14.3193 | 24.2899 |
| | A264471 | 20.8 | 20.2 | 18.7221 | 89.3765 | 17.3907 | 34.3833 | 16.9579 | 22.4559 |
| AM | A0112 | 18.2 | 17.2 | 14.6463 | 114.981 | 13.0098 | 42.784 | 12.7043 | 22.3061 |
| | A0114 | 22.8 | 22 | 18.915 | 104.6125 | 17.3287 | 37.9736 | 17.0273 | 21.8626 |
| | A0115 | 18.2 | 17.2 | 14.6554 | 101.4683 | 13.2178 | 36.1459 | 12.7362 | 19.7837 |
| | A4603 | 20 | 19.4 | 16.49 | 114.2773 | 14.8905 | 38.5233 | 14.4713 | 21.3979 |
| | A4604 | 15.2 | 14.8 | 12.8591 | 126.1443 | 11.3125 | 45.748 | 11.0679 | 23.6763 |
| | A4606 | 16.4 | 16 | 13.4001 | 98.5016 | 11.9592 | 30.612 | 11.7111 | 19.2074 |
| | A4607 | 17.4 | 17 | 14.3709 | 112.1172 | 12.8265 | 35.58 | 12.5487 | 20.816 |
| | A4608 | 26.4 | 24 | 17.9614 | 153.6219 | 15.5811 | 44.2128 | 15.0876 | 21.6128 |
| | A4610 | 17.6 | 17 | 14.2914 | 154.0306 | 12.3834 | 52.4925 | 12.0575 | 24.994 |
| | A4611 | 17.4 | 16.8 | 14.6177 | 83.9938 | 13.3907 | 29.3374 | 13.0823 | 18.1732 |
| | A4612 | 25.8 | 25.2 | 22.1616 | 223.3902 | 19.619 | 83.0229 | 19.1242 | 34.1177 |
| | A4615 | 23 | 22 | 19.4626 | 208.6474 | 17.091 | 79.6296 | 16.5243 | 31.7376 |
| | A4616 | 20.4 | 19.8 | 16.5888 | 130.7014 | 14.7485 | 43.9226 | 14.3884 | 23.5952 |
| | A4617 | 30.8 | 30 | 25.3631 | 146.6001 | 23.1614 | 48.7894 | 22.7786 | 26.2789 |
| | A4618 | 19.4 | 18.8 | 15.6801 | 99.7054 | 14.0939 | 29.6865 | 13.8598 | 19.2013 |
| | A4619 | 18 | 17 | 14.219 | 93.6479 | 12.8181 | 32.9992 | 12.466 | 18.845 |
| | A4621 | 19.2 | 18.6 | 16.3649 | 151.9507 | 14.4012 | 54.5784 | 14.0542 | 26.3588 |
| | A4624 | 21.4 | 20.6 | 18.3076 | 173.2037 | 16.1946 | 66.8696 | 15.7136 | 29.333 |
| | A4625 | 18.2 | 17.2 | 14.4737 | 105.6332 | 12.9495 | 37.4444 | 12.6901 | 20.6241 |

| | | | | | | | | | |
|---|---|---|---|---|---|---|---|---|---|
| | A4626 | 18.8 | 18.2 | 16.8638 | 204.0196 | 14.7585 | 79.7802 | 14.3101 | 33.9884 |
| PV | A7701 | 14.4 | 14.2 | 13.219 | 169.8755 | 11.4968 | 59.0683 | 11.2722 | 29.9725 |
| | A7705 | 39.2 | 38.6 | 35.4759 | 360.9727 | 31.3317 | 136.2582 | 30.5111 | 53.5343 |
| | A7706 | 37 | 36.4 | 33.5019 | 320.1543 | 29.7575 | 119.1642 | 28.9614 | 48.3125 |
| | A7707 | 22.6 | 22.2 | 20.4039 | 140.3169 | 18.642 | 52.2106 | 18.2631 | 30.4043 |
| | A7708 | 44.6 | 43.8 | 40.4082 | 571.5701 | 34.7183 | 202.4306 | 33.755 | 67.1841 |
| | A7709 | 27.6 | 27.2 | 25.1146 | 371.9118 | 21.4808 | 135.8366 | 20.8081 | 50.2842 |
| | A7710 | 25 | 24.4 | 22.5732 | 334.3563 | 19.3218 | 124.841 | 18.7265 | 45.8426 |
| | A7714 | 24.4 | 23.6 | 21.8221 | 162.6745 | 19.8524 | 63.8837 | 19.4013 | 32.4425 |
| | A7715 | 25.8 | 25.4 | 23.47 | 208.1935 | 21.0413 | 76.7619 | 20.5584 | 35.776 |
| | A7717 | 23.8 | 23.2 | 21.4537 | 311.1687 | 18.36 | 115.9904 | 17.8494 | 44.3945 |
